# Uracil/guanine mismatches trigger MutS HOMOLOG1-dependent mitochondrial DNA double-strand breaks in Arabidopsis

**DOI:** 10.64898/2026.06.10.731485

**Authors:** Chang Zhou, Alejandro Peñafiel-Ayala, Chuande Wang, Daniel B. Sloan, Luis G. Brieba, Shin-ichi Arimura

## Abstract

Angiosperm mitochondrial genomes exhibit exceptionally low nucleotide substitution rates, likely supported by active homologous recombination-mediated repair, while maintaining genome integrity by suppressing illegitimate recombination between imperfectly matched sequences that could otherwise cause deleterious structural rearrangements. The nuclear-encoded MutS HOMOLOG1 (MSH1) protein has been proposed to help resolve this paradox by recognizing mismatches and promoting DNA double-strand break-associated repair responses that suppress both point mutations and illegitimate recombination. However, direct experimental evidence for this proposed activity of MSH1 has been lacking. Here, we demonstrate that targeted base editing using a mitochondrial TALE-cytidine deaminase fusion (mitoTALECD) in Arabidopsis mitochondria induces deletions at the redundant gene *atp6-2*. These deletions phenocopy those generated by mitochondria-targeted TALEN (mitoTALEN) cleavage at the same locus. Notably, deletion events that encompass the base-editing target site were markedly reduced in *msh1* mutant backgrounds, indicating that functional MSH1 promotes their formation. *In vitro* assays further demonstrated that recombinantly expressed MSH1 efficiently recognizes U:G mismatches derived from deoxycytidine-to-deoxyuridine deamination events and performs coordinated double incisions, thereby generating staggered dsDNA breaks with short 3′ overhangs. Together, these findings provide direct experimental support for the long-postulated role of MSH1: the induction of mismatch-triggered DNA double-strand breaks. This MSH1 activity likely promotes the repair of mismatched bases and prevents illegitimate recombination by rejecting annealing between imperfect repeat sequences, thereby helping maintain the characteristic mutational stasis and recombinational dynamics of angiosperm mitochondrial genomes.

## Main

Mitochondria are essential organelles that support eukaryotic life through ATP production, metabolic integration, and the regulation of programmed cell death. They possess their own genome (hereafter, the mitogenome), which encodes key components of the respiratory machinery and remains fundamental to mitochondrial function (Morley and Nielsen 2017). In flowering plants, mitochondrial DNA (mtDNA) is notable for its highly variable and often exceptionally large size—ranging from less than 100 kb to over 10 Mb—and for undergoing frequent structural rearrangements. These dynamic features coexist with remarkably low nucleotide substitution rates in gene-coding regions, creating a striking contrast between structural fluidity and mutational stasis (Kubo and Newton 2008, Gualberto, Mileshina et al. 2014, Chevigny, Schatz-Daas et al. 2020). Unlike the nuclear genome, which primarily repairs double-strand breaks (DSBs) through non-homologous end joining (NHEJ), plant mitochondria rely predominantly on homologous recombination (HR) for DSB repair (Gualberto, Mileshina et al. 2014). The multicopy nature of the mitogenome inside cells (Preuten et al., 2010) and frequent fission and fusion of plant mitochondria (Arimura 2004) provides templates for HR, enabling accurate repair of DSBs. At the same time, the abundance of dispersed repeats drives structural rearrangements: large repeats (>1 kb) with perfect identity recombine readily, whereas recombination between intermediate (50–500 bp) or short (<50 bp) repeats, typically sharing only partial sequence identity, is strongly suppressed. This suppression is mediated by the nuclear-encoded surveillance factor MutS HOMOLOG1 (MSH1), which represents an enigmatic lineage within the larger MutS gene family and acts as a key guardian of organellar genome stability (Abdelnoor, Yule et al. 2003, Lin, Nei et al. 2007, Ogata, Ray et al. 2011, Sloan, Broz et al. 2025). Genetic studies have shown that MSH1 plays a central role in suppressing illegitimate recombination between non-identical homologous sequences (Martinez-Zapater, Gil et al. 1992, Abdelnoor, Christensen et al. 2006, Davila, Arrieta-Montiel et al. 2011, Odahara and Sekine 2018, Odahara 2020, Zou, Zhu et al. 2022), and also contributes to the exceptionally low mutation rates observed in plant mitochondria and plastids (Wu, Waneka et al. 2020, Broz, Hodous et al. 2025, Lencina, Prina et al. 2025). These two defining features—suppression of illegitimate recombination and maintenance of extremely low organellar mutation rates—are thought to arise from the dual activities of MSH1: a mismatch-recognition MutS domain and a GIY–YIG endonuclease domain that is located at its C-terminus and distinguishes this protein from the rest of the MutS family. These structural features have led to the hypothesis that MSH1 recognizes mismatches and introduces a DSB at the site, thereby removing the mutated DNA strand and preventing annealing between imperfectly matched repeat sequences. This mismatch-triggered cleavage is proposed to channel repair through high-fidelity HR using an intact template, ultimately ensuring both accurate mutation correction and suppression of illegitimate recombination **(Extended Data Fig. 1a)** (Christensen 2014, Ayala-Garcia, Baruch-Torres et al. 2018, Broz, Keene et al. 2022). However, the central premise of this hypothesis that plant MSH1 mediates mismatch-dependent DNA cleavage has not yet been demonstrated directly (Christensen 2014). Recent studies have shown that the GIY–YIG domain of recombinant *Arabidopsis* MSH1 binds to branched DNA structures (Fukui, Harada et al. 2018) and that the full-length recombinant protein exhibits endonuclease activity toward recombination intermediates such as displacement loops (Penafiel-Ayala, Peralta-Castro et al. 2024). Nevertheless, the key prediction of the model, mismatch-triggered DNA cleavage by plant MSH1, remains unproven. Here, we report the discovery of deletion events occurring precisely at the site of targeted base editing and demonstrate through genetic analyses that their formation depends on MSH1. We further show that purified MSH1 can introduce dsDNA breaks at mismatched sites within the same DNA sequence substrates *in vitro*, explaining the long unresolved question of how MSH1 executes genome maintenance.

## Results

### Occasional DSBs observed by mitochondrial-targeted base editing

While employing a mitochondrial base editor, mitochondria-targeted transcription activator-like effector–cytidine deaminases (mitoTALECDs), to induce C-to-U conversions at the upstream region of ATP synthase-related genes, we observed a loss of PCR amplification at *atp6-2* in several T□ plants. This phenotypic outcome was unexpected, as cytidine deaminases function by mediating single-nucleotide chemical conversions without cleaving the DNA phosphodiester backbone, a process typically expected to yield C-to-T substitution rather than genomic deletions. Loss of *atp6-2* was shown to have no effect on growth because of the presence of a redundant gene, *atp6-1,* in *A. thaliana* ecotype Columbia-0 (Col-0) **(Extended Data Fig. 2a)** (Arimura, Ayabe et al. 2020). The lack of PCR amplification for *atp6-2* is similar to our previous observations using mitochondria-targeted transcription activator-like effector nucleases (mitoTALENs) (Kazama, Okuno et al. 2019, Arimura, Ayabe et al. 2020), which induce DSBs at specific mitochondrial loci via a FokI nuclease domain. mitoTALECD consists of DNA-binding domain TALE arrays, a double-stranded DNA-targeting cytidine deaminase DddA_tox_ (CD), and a uracil glycosylase inhibitor (UGI), but lacks any endonuclease or nickase domain or activity (Mok, de Moraes et al. 2020) **(Fig. 1a)**. Given this design, we hypothesized that base editing–induced U:G mismatches might trigger MSH1-dependent DNA cleavage (Christensen 2014).

**Fig. 1:**
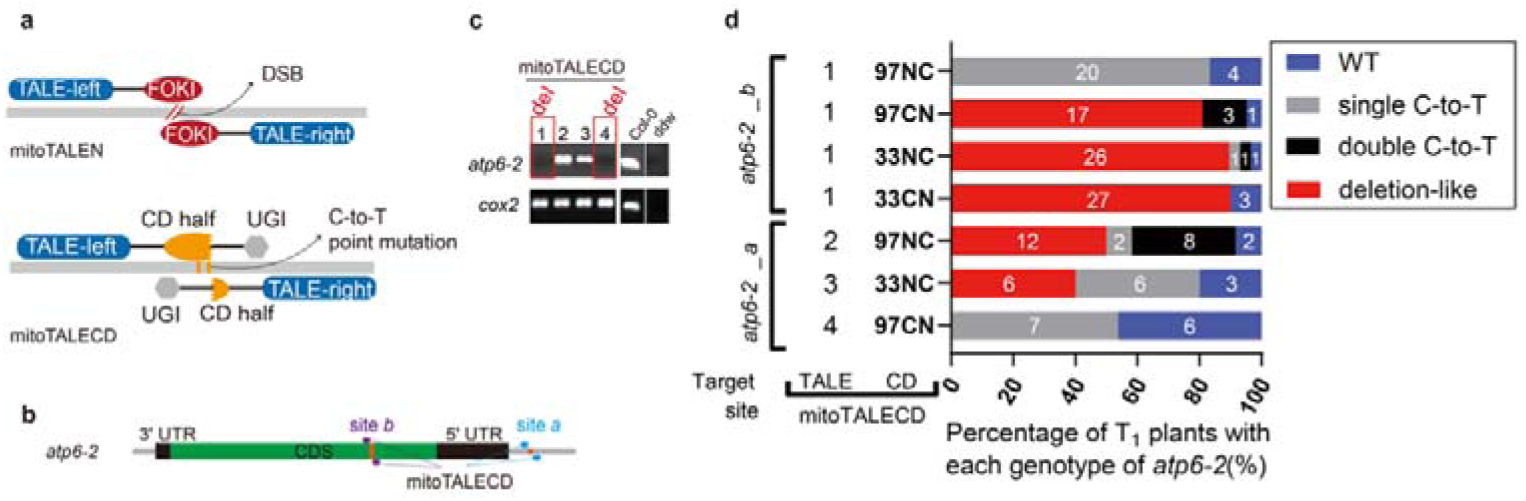
C-to-T base editing at *atp6-2* induces deletion-like genotypes. **a,** Schematic representation of mitoTALEN and mitoTALECD architectures. FokI, FokI restriction endonuclease cleavage domain DSB, double-strand break; CD, cytidine deaminase DddA_tox_; UGI, uracil glycosylase inhibitor. **b,** Location of two target sites, *atp6-2_a* and *atp6-2_b*. Further details, including mitoTALECD variants, target site sequences of *atp6-2*, and the positional relationship of *atp6-2* with *atp6-1* and the large repeat 1-1 and 1-2, are provided in Extended Data Fig. 2a. **c,** PCR genotyping of representative T□ plants transformed with *atp6-2_a*_mitoTALECD_G1333NC. Lines failing to amplify the *atp6-2* locus are highlighted in red boxes. Amplification of *cox2* served as a positive control. **d,** Distribution of *atp6-2* genotypes in T□ plants transformed with different combinations of TALE repeats and cytidine deaminases at sites *a* and *b*. The Y-axis indicates TALECD variants, and the X-axis shows the percentage of each genotype category (wild type, single or double C-to-T edits, and deletion-like events). n = number of T□ plants analyzed.

### mitoTALECD-induced deletions in two cases at the sites of *atp6-2*

In base editing experiments targeting the upstream region (referred to as site *a*) of a*tp6-2* **(Fig. 1b)**, we observed a loss of PCR amplification of a 410-bp fragment flanking the targeted site in several T□ plants edited with two different versions of mitoTALECD (G1333NC and G1397NC) that differ in both their cytidine deaminase domains and TALE array configurations **(Fig. 1c**, **1d**, **Extended Data Fig. 2b**, **3a**; see Supplementary Table 1 for details). In contrast, a third mitoTALECD (G1397CN) targeting this site did not result in losses of PCR amplification and instead produced only C-to-T substitutions (albeit at a relatively low frequency of less than 50%; **Fig. 1c**). To test the reproducibility of this phenomenon, we designed four additional mitoTALECDs targeting a specific coding region of *atp6-2* (referred to as site *b*; **Fig. 1b, Extended Data Fig. 2a**). Genotyping of T□ plants revealed deletion patterns with three of these constructs (87%, 90%, and 91% of T□ individuals), whereas only C-to-T substitutions were observed with the fourth (G1397NC) construct **(Fig. 1d, Extended Data Fig. 3b).** We also verified the absence of any construct mix-up by performing PCR to detect the presence of CD (for mitoTALECD) and FokI (for mitoTALEN) coding regions in these lines **(Extended Data Fig. 3c)**.

When sequencing the amplified PCR fragments of *atp6-2*, we observed that the G1397NC base editor targeting the *b* site predominantly introduced a single C-to-T substitution at the 10^th^ C (C10, hereafter) in between the TALE-recognized sequences, whereas G1333NC and G1397CN induced double substitutions at this site (C10&G13 and G5&C10, respectively) in some of the small number of their T_1_ plants that did not carry deletions **(Fig. 1d, Extended Data Table 1)**. Overall, five of the seven mitoTALECD constructs that successfully edited site *a* or *b* of *atp6-2* caused deletion genotypes in their treated plants **(Fig. 1d)**. The heritability of the deletion or non-deletion was confirmed in the T_2_ progenies of three T_1_ lines each at the *atp6-2_b* site (TALECD_G1397CN) **(Extended Data Fig. 4)**.

To verify the structure of the recombined mitogenomes, we performed Illumina short-read sequencing on six T□ null segregants lacking the nuclear T-DNA expression cassette of the base editor *atp6-2_b*_mitoTALECD_G1397CN. Short-read mapping revealed ∼270 bp deletions in four T□ lines and ∼450 bp deletions in two lines **(Extended Data Fig. 5)**, corresponding to two types of mitogenome rearrangements: (i) deletions between a pair of direct repeats (named Y2 and Y3 [(Davila, Arrieta-Montiel et al. 2011, Zou, Zhu et al. 2022)]) caused by HR that excises the intervening *atp6-2* sequence, and (ii) deletions replaced by a duplicated *atp6-1* region generated through recombination between the V repeat and a long repeat **(Extended Data Fig. 6)**. Together, these results demonstrate that C-to-T editing at *atp6-2_b* induces DSBs that are subsequently repaired by HR, producing at least two distinct mitogenome structures.

In summary, C-to-T base editing at two *atp6-2* sites using five distinct mitoTALECDs generated deletion genotypes in *Arabidopsis thaliana* Col-0. Illumina sequencing of six T□ null segregants edited at *atp6-2_b* confirmed the presence of homoplasmic deletions and revealed two distinct HR outcomes, designated as Y2/3 and V1/3 recombination types. Notably, the Y2/3 recombination type was identical to the rearrangements previously observed in mitoTALEN-targeted *atp6-2_b* plants that repair and bypass the edited region (Arimura, Ayabe et al. 2020), suggesting that mitoTALECD-induced deletions likely originate from DSBs occurring near the target site.

### Deletion at *atp6-2_b* correlates with the presence of MSH1

The unexpected observation of *atp6-2* deletions following C-to-T base editing prompted us to test whether MSH1 contributes to their occurrence. We introduced the same mitoTALECD in the *msh1* mutant background, *msh1-2* (i.e. *chm1-2*) (Abdelnoor, Yule et al. 2003). Because *msh1* mutants have already accumulated structural mutations in their mitogenomes, *msh1* individuals retaining *atp6-1*, *atp6-2*, and *cox2* were selected by multiplex PCR to prevent using *msh1* individuals spontaneously lacking *atp6-1* or *atp6-2* **(Fig. 2a)**. The presence of *atp6-1* is necessary to avoid lethality upon deletion of *atp6-2*. Selected *msh1* lines were then transformed with the mitoTALECD *atp6-2*_*b*_G1397CN. Genotyping of the resulting T_1_ transformants revealed a stark contrast between wild-type (*MSH1/MSH1*) and *msh1-2* mutant (*msh1/msh1*) backgrounds. In the wild-type background, 81% of T_1_ lines exhibited deletions at *atp6-2*_*b*, whereas 9.7% of *msh1-2* lines displayed such deletions, indicating a correlation between the presence of functional MSH1 and deletion frequency **(Fig. 2a)**. To confirm this correlation under a uniform and intact mitogenome background, we generated heterozygous

**Fig.2:**
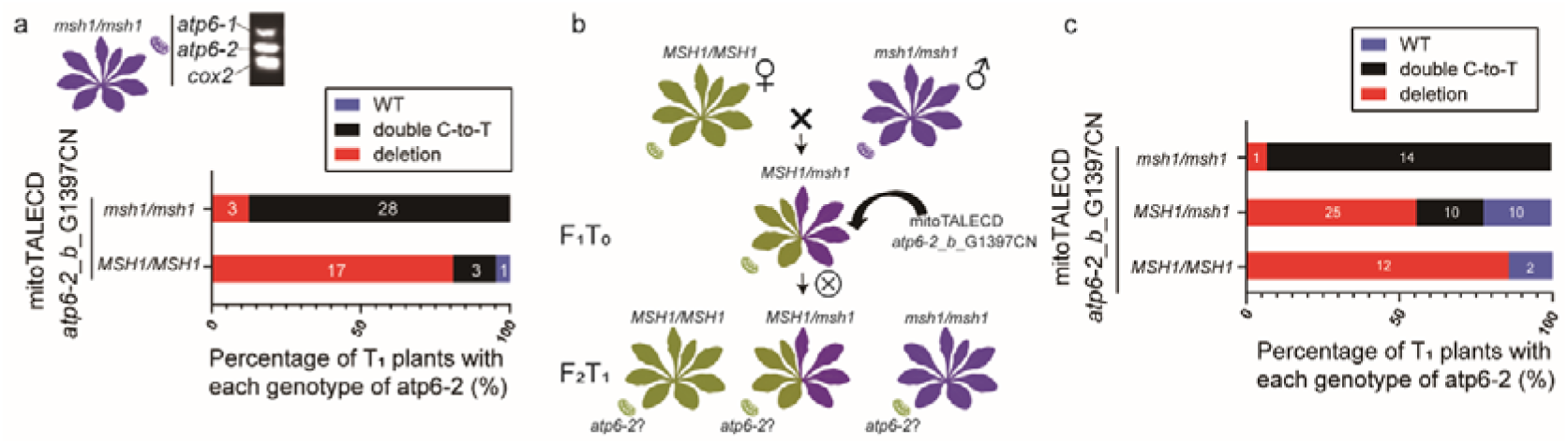
Deletion rate at *atp6-2_b* is strongly associated with nuclear *MSH1* genotype. **a,** Distribution of *atp6-2* genotypes in T_1_ plants transformed with mitoTALECD *atp6-2_b*_G1397CN in wild type (*MSH1/MSH1*) and *msh1* mutants (*msh1/msh1*). *msh1* plants that retained *atp6-1*, *atp6-2* and *cox2* were selected for transformation. The numbers within the bars indicate the number of corresponding plants. **b,** Crossing design to test the effects of MSH1 genotype on rates of deletion at *atp6-2_b* under a wild-type mitochondrial background. Using a wild-type maternal plant (green) and a homozygous *msh1-2* mutant (purple) pollen donor, we generated heterozygous *msh1-2* F_1_ individuals that carried cytoplasmic genomes inherited from a wild-type lineage (as indicated by the green mitochondrion). Heterozygous *msh1* F_1_ individuals were then transformed with mitoTALECD *atp6-2_b*_G1397CN as T_0_ plants using the floral dipping method. After selfing the F_1_T_0_, we genotyped the resulting F_2_T_1_ progeny to identify homozygous *msh1* (*msh1/msh1*) mutants, heterozygous *msh1* (*MSH1/msh1*) mutants and homozygous wild-type (*MSH1/MSH1*) individuals. All F_2_T_1_ were then genotyped for *atp6-2_b*. **c,** Distribution of *atp6-2* genotypes in F_2_T_1_ plants transformed with mitoTALECD *atp6-2_b*_G1397CN. The Y-axis represents the three *MSH1* genotypes of F_2_T_1_, and the X-axis represents the percentage of different genotypes of *atp6-2_b*. The numbers within the bars indicate the number of corresponding plants.

*MSH1/msh1* F_1_ individuals by crossing Col-0 (♀) with *msh1-2* (♂), thereby ensuring presence of the wild-type intact mitogenome due to maternal inheritance **(Fig. 2b)**. These F_1_ plants were transformed with *atp6-2*_*b*_G1397CN, and their self-crossed F_2_ progeny (F_2_T_1_) were genotyped for both *MSH1* **(Extended Data Fig. 7)** and *atp6-2*_*b* **(Extended Data Fig. 8)**. Among 74 F_2_T_1_ plants with confirmed T-DNA and intact mitochondrial background, deletions at *atp6-2*_b were detected in 86% of *MSH1/MSH1* individuals, 56% of *MSH1/msh1* heterozygotes, and 7% of homozygous *msh1-2* mutants **(Fig. 2c)**. These results provide genetic evidence that MSH1 facilitates the formation of deletions at *atp6-2*_*b* following base editing.

### MSH1 cleaves mitochondrial-targeted base editing sites

To test if MSH1 promotes deletions at the *atp6-2* locus by cleaving U:G mismatches generated by cytidine deamination, we assayed the *atp6-2*_*a* and *atp6-2*_*b* editing sites using fluorescently labeled dsDNA (Cy5 on one strand and FAM on the other) and purified full-length MSH1 protein *in vitro* (Peñafiel-Ayala, Zhou et al. 2026). For *atp6-2b*, we followed the cytidine deamination pathway by synthesizing DNA substrates with modifications to the C:G base pair corresponding to the targeted C10 (10^th^ nt from the left TALE binding site) to generate three editing states: a U:G mismatch, a U:A pair, and a fixed T:A mutation (**Fig. 3a**). Enzymatic reactions run on a denaturing gel show that MSH1 cleaves the uracil-containing oligonucleotide in the substrate containing the U:G mismatch but exhibits minimal cleavage for U:A and T:A pairs and no cleavage for the C:G wild-type sequence (**Fig. 3b, lanes 13 to 20**). Similarly, MSH1 efficiently cleaves the complementary oligonucleotide on a U:G mismatch, U:A and T:A pairs, but not on the G:C wild-type sequence (**Fig. 3b, 1 to 8**). The main cleavage sites are located 12 nt from the deamination site C10 in the 5’ direction yielding a 20-nt fragment from the 5′ end of the 60-mer labeled strand (this product is highlighted with a cyan triangle) and 15 nt in the 3′ direction of the complementary strand of residue C10 yielding a 43-nt fragment from the 5′ end of the complementary 60-mer labeled oligonucleotide (this product is marked with a cyan triangle). The cleavage positions create a DSB (**Extended Data Fig. 9a**), resulting in two dsDNA fragments: a 17-bp Cy5-labeled fragment and a 40-bp FAM-labeled fragment, both bearing 3-nt 3′ overhangs (**Fig. 3c**). We further analyzed two additional substrates based on the outcomes of *atp6-2_b* mitoTALECD_G1333NC, which induced double substitutions at C10 and G13 and a high frequency of deletion-like events (89.7%) in T_1_ plants **(Fig. 1d, Extended Data Fig. 3b)**. These substrates, featuring deamination at the G13-paired cytosine either alone (G:U) or with the C10 mismatch (U:G), were tested to determine if such clustered lesions potentiate MSH1-directed cleavage **(Fig. 3a)**. In both cases, the 20-nt and 43-nt cleavage products with respect to the 5’-end of the labeled oligonucleotides were observed following MSH1 incubation (**Fig. 3b, lanes 21-24 and 9-12**). However, both substrates produce additional cleavage products of 28 nt (C10/G13 strand) and 34 nt (complementary strand) (**Fig. 3c, lanes 30 and 32, lanes 26 and 28,** green triangles and Fig 5S). These cleavage products localize 7 and 9 nt from the G13 mismatch, generating a new DSB with a 2-nt staggered end. Accordingly, the reaction yields dsDNA fragments of 32 bp and 26 bp, detected in the FAM and Cy5 channels, respectively (Fig. 3c, lanes 30 and 26; green labels).

**Fig. 3:**
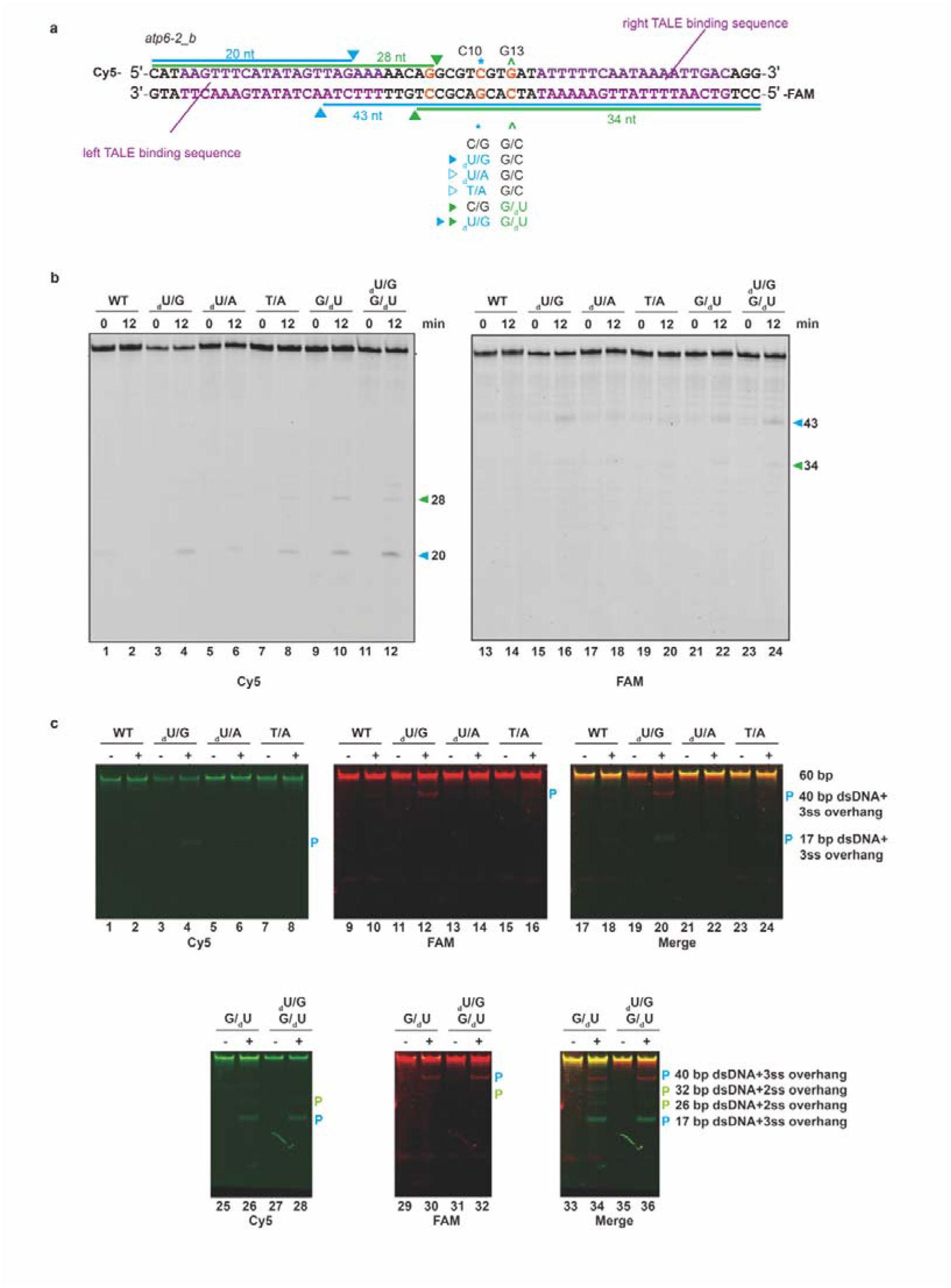
MSH1 cleaves *atp6-2_*b sequences processed by mitoTALE cytidine deaminases. **a,** Schematic representation of the assembled dsDNA corresponding to the *atp6-2_*b target site. TALE binding sites are highlighted in purple, * and symbols denote the mutated sites at the10^th^ C and 13^th^ G downstream from the left TALE binding site, respectively. The modified cytidines are labeled in blue and pale green, and the cleavage products are indicated by triangles. **b,** 12% denaturing PAGE showing cleavage of 5′-Cy5 and 5’-FAM labeled substrates containing variants in the *atp6-2_*b sequence (C:G wild-type, deaminated cytidine to uracil, U:A pair, and T:A fixed mutation). MSH1 cleaves the U:G mismatch at position 43-bp from the 5’ of the oligonucleotide complementary to the lesion and exhibits minimal cleavage for U:A and T:A pairs. MSH1 cleaves 20-bp from the 5’ of the oligonucleotide harboring a U:G mismatch, U:A and T:A pairs, but not on the G:C wild-type sequence (cyan triangle). The introduction of the double U:G pair creates an alternative cleavage site colored in pale green. **c,** Native polyacrylamide gel showing dsDNA cleavage at the *atp6-2_*b sequence. MSH1 mainly cleaves the substrate with the U:G mismatch at a defined position generating a FAM (red) and a Cy-5 (green) labeled dsDNA products (lanes 4, 12, and 20; annotated with a cyan “P”). The substrate with a double U:G mismatch (at C10 and G13), new cleavage products are observed in concordance with the U:G mismatch at G13. The 32-bp FAM-labeled product is visualized as a red band (lanes 26 and 28), whereas the 26-bp Cy5-labeled product appears as a green band (lanes 30 and 32). Notably, the 32-bp FAM-labeled product co-migrates with the unpaired Cy5-labeled oligonucleotide, complicating its resolution in the merged channels.

Each fragment carries the expected complementary 2-nt 3′ overhang, consistent with offset cleavage on opposite strands at the defined positions flanking G13. In the merged channel, a non-hybridized ssDNA species masks the ∼32-nt FAM-labeled dsDNA product (**Fig. 3c**, lanes 33 to 36). This cleavage pattern suggests that MSH1 recognizes the newly deaminated U:G base pair as a mismatch and generates new dsDNA products. These MSH1 cleavage products are potential substrates for exonucleases to initiate RecA-mediated HR (Gualberto, Mileshina et al. 2014, Gualberto and Newton 2017).

For *atp6-2*_a, we also followed the cytidine deamination pathway for C12, the 12^th^ nucleotide from the left TALE binding site (**Fig. 4a**). In contrast to wild-type *atp6-2*_b sequence, which is barely cleaved, MSH1 exhibits potent DNA cleavage activity on the wild-type *atp6-2*_a sequence, producing incisions located 37 nt from the 5′ end of the editing strand and 26 nt from the 5′ end of the complementary strand (**Fig. 4b, lane 14 and 2, Extended Data Fig. 9b**). This non-specific cleavage results in two dsDNA products of 23 and 34 nt labeled with FAM and Cy-5, respectively (**Fig. 4c; labeled as “off”**). This promiscuous dsDNA cleavage is also observed in substrates in which the C:G pair is modified to a U:G mismatch, a U:A pair, and a T:A fixed mutation (**Fig. 4b**). In addition, MSH1 introduces U:G mismatch-dependent incisions 20 nt from the 5′ end in the deaminated strand and 42 nt from the 5′ end of the complementary strand, leaving dsDNA products of 18 bp and 40 bp respectively, with 2-nt 3′ overhangs. (**Fig. 4c**). Those incisions match the cleavage pattern observed in dsDNA substrates with designed sequence and in sequences harboring the wild-type *atp6-2*_*b* sequence with a U:G mismatch (Peñafiel-Ayala, Zhou et al. 2026), with the exception that the overhang is 2 nt instead of 3 nt (**Fig. 4a**). Interestingly, on this substrate, cleavage at non-specific sites is more pronounced than at the mismatch-induced sites, exceeding the intensity of the bands typically observed for mismatch-containing templates or the *atp6-2*_a sequence harboring a U:G mismatch. The promiscuous activity of MSH1 correlates with its relaxed specificity observed in the presence of Mn^2+^ instead of Mg^2+^ (Peñafiel-Ayala, Zhou et al. 2026) (**Fig. 4a**).

**Fig. 4.**
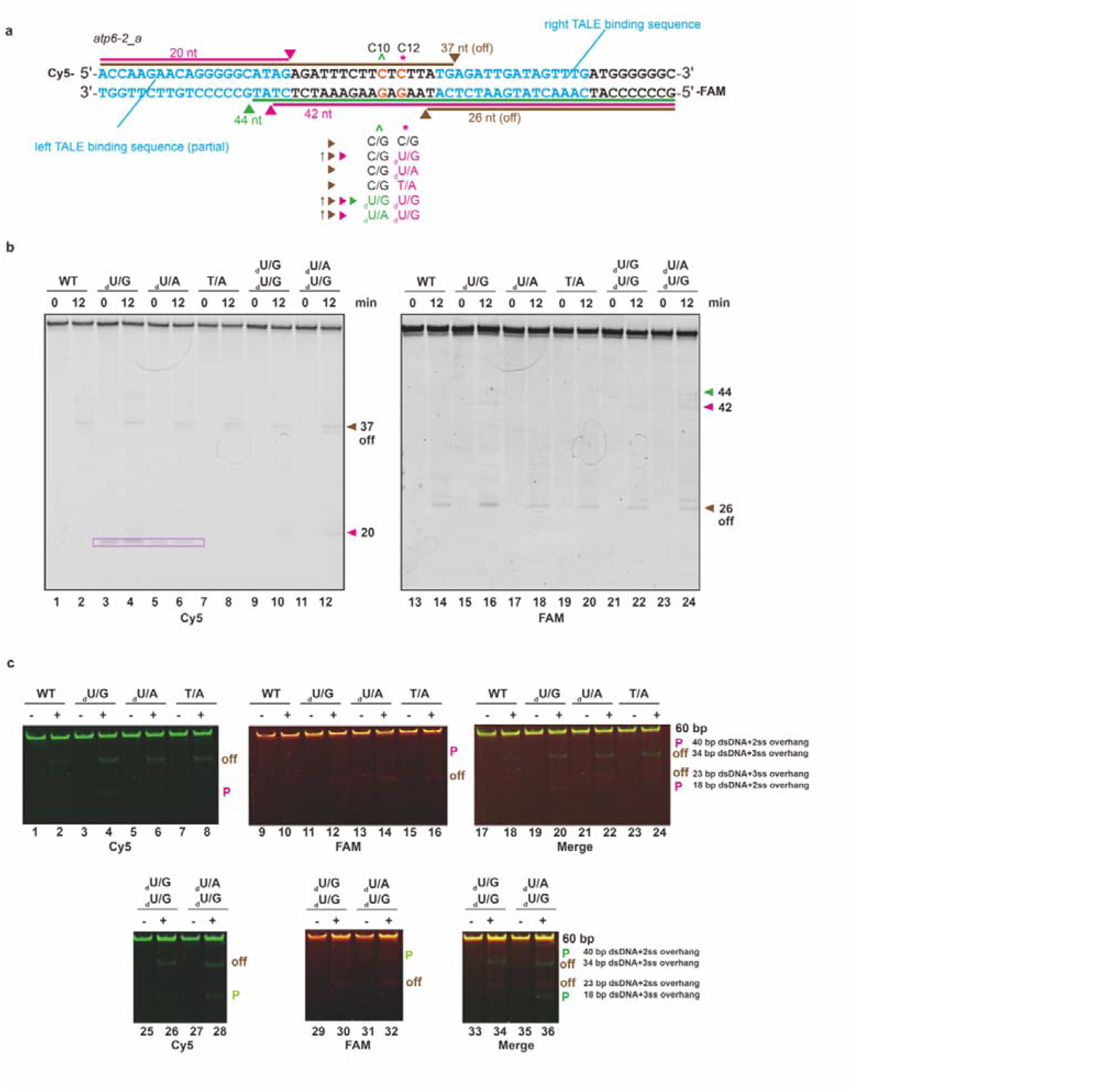
MSH1 cleaves *atp6-2*_a sequences processed by mitoTALE cytidine deaminases in a mismatch-dependent and in a mismatch-independent manner. **a,** Schematic representation of the dsDNA corresponding to the *atp6-2*_a target site. TALE binding sites are highlighted in cyan, and the and * symbols denote the C10 and C12 mutated sites downstream from the left TALE binding site. **b,** 12% denaturing PAGE showing cleavage products of 5′-Cy5 and 5’-FAM labeled substrates containing variants in the *atp6-2*_a sequence (C:G wild-type, deaminated cytidine to uracil, U:A pair, and T:A fixed mutation). The positions of the non-specific MSH1 cleaving products of 26 and 37 nt are indicated by brown triangles. The mismatch-specific cleavage sites of 20 and 42 nt are indicated by pink triangles. Green triangles indicate the cleavage site potentiated by the U:G mismatch at C12. The purple rectangle denotes a contaminating oligonucleotide fragment (lanes 15 to 18). **c,** Native polyacrylamide gel showing dsDNA cleavage at the *atp6-2_*a sequence. The non-specific cleavage products are labeled “off,” and products corresponding to mismatch-dependent recognition are labeled “P.”

To further assess MSH1 cleavage at the *atp6-2_a* sequence, we designed two additional substrates in which both edited cytidines, C10 and C12, were converted to uracil. In these constructs, C12 position always forms a U:G mismatch, whereas C10 position either forms a U:G mismatch or pairs with adenine (U:A). In both substrates, MSH1 maintains cleavage at the promiscuous site (**Fig. 4b, lanes 21–24 and 9–12**); however, the overall intensity of mismatch-dependent cleavage increases, and four distinct dsDNA products become clearly resolved on a native agarose gel (**Fig. 4c, lanes 25–36**). Two of these products correspond to non-specific (off-target) DSBs, while the remaining two are dependent on the presence of U:G mismatches. Notably, introducing a second U:G mismatch at C10, located two nucleotides from the original mismatch at C12, generates an additional cleavage product of 44 nt—2 nt longer than the original 42-nt fragment—consistent with restoration of a 4-nt staggered overhang (**Fig. 4b, lane 24**). Deamination at C10 enhances the cleavage efficiency of MSH1, indicating that closely spaced U:G mismatches may potentiate its nuclease activity. The identity of uracil in the DNA substrates was verified by UDG-mediated excision followed by strand cleavage on a denaturing urea gel, where only uracil-containing substrates produced defined fragmentation patterns (**Extended Data Fig. 10)**. Overall, these results indicate that MSH1 can recognize and cleave DNA sequences in a promiscuous manner, exhibiting relaxed specificity that enables both mismatch-dependent and non-specific DNA cleavage.

## Discussion

In this study, we show that targeted C-to-T base editing at the mitochondrial *atp6-2* locus induces MSH1-dependent deletion and recombination via flanking repeat sequences, resulting in a structurally altered mitogenome that lacks the *atp6-2* locus. This process is likely initiated by the recognition and cleavage of a U:G mismatch intermediate, generated by targeted base editing by mitoTALECD. These findings provide both *in vivo* and *in vitro* evidence for the mismatch-directed cleavage activity of MSH1 in plant mitochondria—an activity that has remained unproven until now.

In plants, mitochondria share their genome via extensive and frequent fusion and fission events (Arimura, Yamamoto et al. 2004, Preuten, Cincu et al. 2010, Wang, Zhang et al. 2010, Arimura 2018). Under natural circumstances, spontaneous damage and mismatches arising on a single mtDNA molecule would be recognized and cleaved by MSH1, followed by accurate repair via HR, probably through a break-induced replication pathway (Gandini, Garcia et al. 2023) using intact copies in the same mitochondrion or another mitochondrion via fusion (Paszkiewicz, Gualberto et al. 2017, Arimura 2018). In contrast, our experimental system would be able to introduce C-to-U conversions at the targeted sites in multiple copies of *atp6-2* through repeated attack by mitoTALECD base editor enzymes. This editing and cleavage of multiple mtDNA copies likely overwhelmed the availability of intact wild-type templates and impeded precise reconnection by homologous repair. Consequently, rarely occurring ectopic recombination via flanking repeat sequences appeared and resulted in deletions of the *atp6-2* locus. These deletions were detected as failed PCR amplification of the target region in T□ plants. The observed PCR failure closely resembled the previously reported pattern in mitoTALEN-based editing experiments(Arimura, Ayabe et al. 2020), supporting the notion that C-to-U conversions activated MSH1-dependent DNA cleavage. These deletions were strongly suppressed in favor of C-to-T mutations in the *msh1* mutant background (**Fig. 2**). It is worth noting that a minor fraction of *atp6-2_b* deletions was still observed in *msh1* mutants with mitoTALECD treatment. This phenotype, though seemingly contradictory to MSH1’s involvement in executing DNA cleavage, is consistent with its established role as a gatekeeper of mitogenome stability. Beyond participating in mismatch-directed break induction, MSH1 actively suppresses non-homologous or ectopic recombination across intermediate alternative repeats (Davila et al., 2011; Zou et al., 2022). The loss of MSH1 function is known to trigger substoichiometric variations and spontaneous rearrangements via flanking genomic repeats.

To our knowledge, such large-scale deletions have not been previously reported in studies employing mitoTALECD to induce C-to-T conversions in plant mitogenomes (Kang, Bae et al. 2021, Nakazato, Okuno et al. 2022, Tamburino, D’Agostino et al. 2025). In addition to the redundancy of *atp6-2,* which enables us to produce knock-out plants without lethality, the local genomic context—characterized by one of the ends having a recombinationally active long repeat of 6.6 kb and another having a high density of intermediate and small repeat sequences likely contribute to the feasible detection of the observed recombination outcomes. In addition, another unusual and fortuitous feature of *atp6-2* is that its upstream region has no identical sequence in the nucleus, as almost all other mitogenome content is captured in a large nuclear-encoded mtDNA sequences (NUMT) on nuclear chromosome 2(Fields, Waneka et al. 2022). As a result, it is easy to detect the absence of PCR due to loss from the mitogenome without confounding effects from NUMT copies (Arimura, Ayabe et al. 2020).

Notably, among the four mitoTALECD constructs targeting *atp6-2_b* in this study, *atp6-2_b_G1397NC* induced single C-to-T conversions with high efficiency, yielding homoplasmic edits in 20 out of 24 T□ plants (83%) without detectable deletions. In contrast, deletions were observed with the other three constructs, two of which*, atp6-2_b*_G1397CN and *atp6-2_b*_G1333NC, produced double C-to-T mutations. These results suggest that MSH1 may differentially recognize distinct spatial configurations of mismatches arising from different types of base editing outcomes. The clustered distribution of mutations in *msh1* mutants (Wu, Waneka et al. 2020, Zou, Zhu et al. 2022) also implies that MSH1 recognition may be context-dependent, guided by local sequence or structural cues.

Moreover, our previous attempt to induce an A-to-G conversion (via A-to-I deamination activity) at the *atp6-2_b* locus using monomeric TALEAD (mTALEAD) did not result in the large-scale deletions observed following C-to-T editing (Zhou, Okuno et al. 2024). However, it is important to note that mTALEAD exhibited relatively low editing efficiency at this site, with the highest heterozygous editing rate at the eighth thymine in T□ plants reaching only 27%. Consequently, only a limited number of copies with A-to-G alleles were generated. The absence of detectable deletions suggests that MSH1 does not efficiently recognize A:I mismatches. Consistent with this observation, *in vitro* assays using MSH1 and dsDNA substrates containing inosine demonstrated that MSH1 fails to cleave A:I and C:I pairs and exhibits only limited cleavage activity toward G:I and T:I mismatches. Moreover, studies employing artificial DNA substrates revealed that U:G mismatches are the preferred substrate for MSH1, being cleaved more efficiently than T:G mismatches (Peñafiel et al., 2025). Notably, Broz *et al*. recently measured *de novo* organellar mutation spectra using *msh1* mutation-accumulation (MA) lines, a system that minimizes selective filtering of new mutations. Their results showed a strongly GC-biased spectrum dominated by AT→GC transitions in plastids, whereas mitochondrial mutations were largely GC-neutral, displaying no obvious bias toward any specific transition class (Broz, Hodous et al. 2025). These findings indicate that, under near-neutral conditions, C→T transitions are not the predominant mutation type in mitochondria. More broadly, the contrasting mutation spectra between plastids and mitochondria suggest that the mutational consequences of MSH1 loss are complex and organelle-specific, rather than restricted to a single mismatch category.

Our *in vivo* results are corroborated by biochemical data that shows that recombinantly expressed MSH1 recognizes and cleaves mismatches and cytidine deamination products in a sequence-dependent manner. MSH1 exhibits enhanced cleavage at U:G mismatches, whereas U:A pairs are processed less efficiently (Peñafiel-Ayala, Zhou et al. 2026), a preference that is consistent with human MSH proteins (Mazurek, Berardini et al. 2002, Larson, Bednarski et al. 2008). On a substrate that mimics the *atp6-2_b* locus, MSH1 recognize the U:G mismatch catalyzing defined incisions several nucleotides from the mismatch generating staggered

DNA ends, so this distal cleavage events could create 3’-single-stranded overhangs suitable for plant organellar RecA-mediated HR (Mueller, Smith et al. 1995, Coulon, Gaillard et al. 2004, Edgell, Derbyshire et al. 2004, Dunin-Horkawicz, Feder et al. 2006, Kisker, Kuper et al. 2013, Gualberto and Newton 2017, Chevigny, Schatz-Daas et al. 2020). In contrast to the U:G mismatch-dependent cleavage on the *atp6-2_b* locus, MSH1 presents strong cleavage activity on the wild-type *atp6-2_a* sequence in the absence of a mismatch, indicating a degree of promiscuous nuclease activity that is dependent on the sequence context and not the mismatch. This non-specific DNA cleavage parallels the promiscuous cleavage activity observed in several nucleases, such as DNase I and restriction enzymes, where sequence context promotes conformational changes that direct non-specific DNA cleavage (Dryden 2001). Besides non-specific DNA cleavage on the *atp6-2_a* sequence, the introduction of a U:G mismatch induces mismatch-specific cuts that add to the non-specific cleavage sites. The ability of MSH1 to generate multiple DNA cleavage sites suggests that it may contribute to the formation of double-strand breaks (DSBs), which could promote DNA deletions or inversions when repaired through microhomology-mediated end joining (MMEJ) or non-homologous end joining (NHEJ).Thus, our data suggest that, in certain DNA sequence or structural contexts, like the one present in the *atp6-2_a* locus, MSH1 cleavage may promote genomic deletions.

Plant mitogenomes exhibit a suite of striking and seemingly paradoxical features, including exceptionally low synonymous substitution rates, extensive structural rearrangements, rapid divergence of noncoding sequences, high frequencies of repeat-mediated recombination, and dramatic variation in genome size. Alan Christensen (Christensen 2013, Christensen 2014) proposed a unifying hypothesis in which these characteristics arise from a plant-specific DNA repair strategy: mismatches, aberrant recombination intermediates, and other lesions are first converted into DSBs, a process likely catalyzed by the GIY-YIG endonuclease domain of MSH1 and then repaired through diverse DSB repair pathways. Accurate, homology-based repair (e.g., gene conversion) preserves coding sequences and explains their extremely low mutation rates, whereas inaccurate modes of DSB repair (such as break-induced replication and microhomology-mediated recombination,) generate the duplications, chimeric junctions, and genome expansions that accumulate in noncoding regions due to weak purifying selection. Our findings provide a crucial experimental validation of Christensen’s hypothesis and suggest that MSH1-initiated DSB formation constitutes a central mechanism linking mismatch recognition to organellar genome stability. They support a repair pathway in which MSH1-directed dsDNA cleavage is coupled to HR, ultimately promoting accurate lesion or mismatch removal via gene conversion. In sum, our findings support the hypothesis that the low rate of mutation accumulation observed in plant organellar genomes results from an MMR-like pathway that enhances replication fidelity (Modrich and Lahue 1996, McCulloch and Kunkel 2008, Christensen 2014, Kunkel and Erie 2015, On and Welch 2021). By integrating precise mitochondrial base editing with genetic disruption of MSH1, our study strengthens the mechanistic foundation of the MSH1 working model and advances our understanding of how DSB repair processes shape the unique mutational landscape and evolutionary dynamics of plant mitogenomes.

## Methods and materials

### Plant materials and growth conditions

*Arabidopsis thaliana* Columbia-0 (Col-0), *msh1-2* mutants (originally referred to as *chm1-2*; renamed as *msh1-2* following the identification of *CHM1* as *MSH1* in (Abdelnoor, Yule et al. 2003), and transgenic lines were grown at 22°C under long-day conditions (16 h light / 8 h dark). Seeds were surface-sterilized and sown on half-strength Murashige and Skoog (½ MS) medium (pH 5.7) containing 2.3 g l□¹ MS Plant Salt Mixture (Wako), 500 mg l□¹ MES, 10 g l□¹ sucrose, 1 ml l□¹ Plant Preservative Mixture (Plant Cell Technology), 1 ml l□¹ Gamborg’s Vitamin Solution (Sigma–Aldrich), and 5 g l□¹ agar. After germination, seedlings were grown for 2–3 weeks before being transferred to Jiffy-7 pots (Jiffy Products International) and maintained in a greenhouse at 22°C under long-day conditions. Plants were subsequently used for Agrobacterium-mediated transformation.

### Crosses between Col-0 and *msh1-2*

Crosses were performed using wild-type *Arabidopsis thaliana* Col-0 (♀) as the maternal parent and *msh1-2* mutant plants (♂) as the paternal parent. Approximately 4-week-old Col-0 plants were selected, and several unopened flower buds were chosen for crossing. Sepals, petals, and stamens were carefully removed from these buds using fine forceps, leaving only the pistils. Healthy stamens from *msh1-2* plants were collected with forceps, and the anthers were gently brushed onto the stigmas of Col-0 pistils. Crossed stigmas were marked by tying red thread at the corresponding branch nodes. Successful pollination was typically indicated by stigma wilting within ∼10 h. To prevent self-pollination, all other buds and developing siliques on Col-0 plants were continuously removed throughout the hybridization period until F□ seeds were harvested. Genomic DNA was extracted from F□ seedlings, and PCR amplification and Sanger sequencing of the *MSH1* locus were performed to verify successful crossing. Only F□ plants heterozygous for *msh1-2* were selected for downstream experiments.

### Vector constructions

The construction of mitoTALECD vectors followed a previously published study (Nakazato, Okuno et al. 2022). MitoTALECDs targeting specific loci were assembled in Ti plasmids using the Platinum Gate TALEN assembly system (Sakuma and Yamamoto 2016) and multisite Gateway cloning (Thermo Fisher Scientific), as described for mitochondria-targeted TALENs in Kazama et al. (2019). The DNA-binding domains were constructed by assembling left and right TALE arrays, each fused to a split half of the cytidine deaminase DddA_tox_ (CD-half) and the uracil glycosylase inhibitor (UGI), respectively. To ensure mitochondrial targeting, the N-terminus of the mitoTALECDs was fused to the mitochondrial targeting peptide (MTP) of the *Arabidopsis* ATPase δ′ subunit (Arimura, Ayabe et al. 2020).

In this study, the constructs for *atp6-2_*a and *atp6-2_*b were generated by assembling the specific TALE arrays into existing destination vectors that already contained the CD-half and UGI coding sequences, previously modified in earlier work (Nakazato, Okuno et al. 2021, Nakazato, Okuno et al. 2022). Both mitoTALECD constructs contained an Oleosin-GFP expression cassette. Seeds containing the T-DNA insertion were identified under a fluorescence microscope based on green fluorescent protein (GFP) signals and were selected for subsequent experiments. F_2_T_1_ Transformants were also checked by PCR amplification of the neomycin phosphotransferase II (*nptII*) gene, which confers kanamycin resistance in plants and bacteria.

The reading frames from the assembled first, second, and third entry vectors were transferred into the Ti plasmid backbone via a multi-site LR recombination reaction using LR Clonase II Plus (Thermo Fisher Scientific). The second entry vector contained an *Arabidopsis* RPS5A promoter, a mitochondria-localization signal, and the *Arabidopsis* heat-shock protein terminator. The Ti plasmid backbone was derived from the Gateway destination vector pK7WG2 (Karimi et al., 2002), in which the *CaMV 35S* promoter was replaced with the *Arabidopsis* RPS5A promoter, and the MTP coding sequence and Ole1pro::Ole1-GFP cassette (from pFAST02; (Shimada, Shimada et al. 2010) were inserted.

### Plant transformation and screening transformants

Col-0 plants were transformed by floral dipping (Clough and Bent 1998) *Agrobacterium tumefaciens* strain C58C1 that harbored one of the transformation vectors described above. Obtained T_1_ seeds were selected by their seed-specific fluorescence of green fluorescent protein (GFP) (Shimada, Shimada et al. 2010). GFP-positive seeds were sown on the 1/2 MS medium (see section on Plant material and growth conditions) further containing 125 mg l^−1^ of claforan. And T_2_ seeds were sowed on 1/2 MS medium.

### Genotyping T_1_, T_2_, and F_2_T_1_ plants

PCRs for Sanger sequencing were performed using KOD One PCR Master Mix (TOYOBO) with extracted DNA from an emerging leaf or cotyledon at 11 DAS with standard protocols. DNA extraction was performed by placing one leaf in 50 µL of the Plant Very Rapid PCR Isolation Buffer (containing 5 mmol g^−1^ EDTA with pH = 8.0 and 0.1 mol g^−1^ Tris HCl with pH = 9.5) after a 15-min treatment at 98 °C. DNA sequences adjacent to the target DNA were amplified with primer sets. All the primers used in this study were designed with NCBI Primer BLAST (https://www.ncbi.nlm.nih.gov/tools/primer-blast/). Purified PCR products were subjected to Sanger sequencing (Eurofins Genomics) to detect substitution of the targeted bases. The data were analyzed with Geneious prime (v.2025.1.2). Total DNAs for Illumina NGS analyses were extracted from mature leaves of Arabidopsis with DNeasy Plant Pro Kit (QIAGEN).

### Illumina Sequencing and Bioinformatic Analysis

For high-throughput genomic analysis, total DNA extracted from individual T□ plants and controls was sequenced on the Illumina NovaSeq 6000 platform. Bioinformatic processing was performed following the methodology previously described by Nakazato et al. (2021). Briefly, raw sequencing reads were preprocessed to remove low-quality bases and adapter sequences using Platanus_trim (v.1.0.7; http://platanus.bio.titech.ac.jp/platanus_trim). The trimmed paired-end reads were subsequently mapped to the Arabidopsis reference mitochondrial genome (GenBank accession: AP000423.1) using BWA (v.0.7.12) executed in single-end mode. Coverage depths and structural variations flanking the atp6-2 locus were analyzed using Geneious Prime (v.2025.1.2) to evaluate mitochondrial genome rearrangements. All raw next-generation sequencing (NGS) data have been deposited in the NCBI Sequence Read Archive (SRA) database under accession number XXXXXX.

### Yeast heterologous expression and purification of MSH1

BCY123 yeast cells were transformed with pSc-MSH1 (wild type or mutant variants) using a lithium acetate/PEG method with salmon sperm DNA and heat shock, followed by selection on selective media. Transformants were expanded in YP medium, shifted to low glucose, and induced with 3% galactose for 16 h. Cells were harvested, lysed in high-salt HEPES buffer, and clarified by centrifugation. Recombinant AtMSH1 was purified from the soluble fraction by Ni²□-affinity chromatography (HisTrap), followed by heparin chromatography and size-exclusion chromatography (S-300) on an ÄKTA system. Purified fractions were concentrated and stored at −80 °C after liquid nitrogen snap-freezing. The full process of expression and purification is described in detail in (Peñafiel-Ayala, Zhou et al. 2026).

### DNA cleavage reactions using synthetic oligonucleotides

Oligonucleoides sequence and Oligonucleotide combination for DNA substrates annealing were listed in Supplementary Table. 3&4. Fluorescently labeled dsDNA substrates were incubated at 30 °C in a cleaving buffer containing 25 mM BisTris propane (pH 7.5), 50 mM potassium glutamate, 5mM MgCl□, 1 mM DTT, 0.1% BSA, and 1 mM ATP. Reactions typically contained 15 nM dsDNA substrate. Prior to separation of products by 12% denaturing PAGE in 1× TBE buffer, reactions were stopped with 2× stop buffer (95% formamide, 5 mM EDTA). Reaction products separated on a native 8% acrylamide gel were stopped with Ficoll-derived buffer. Reactions were visualized on an Amersham Typhoon™ scanner (Cytiva) with Cy2 or Cy5 filters.

### UDG reactions using synthetic oligonucleotides

Fluorescently labeled oligonucleotides sequences used for MSH1 cleavage were annealed and assayed using UDG (New England Biolabs) to assess the presence of deoxyuridine in the sequences. Substrates were incubated at 37°C using 1x UDG buffer (New England Biolabs), 15 nM dsDNA and 0.5 unit of UDG (New England Biolabs) for 15 minutes. Reactions were stopped with 2× stop buffer (95% formamide, 5 mM EDTA). Products were separated in a 12% denaturing PAGE in 1x TBE buffer. Reactions were visualized on an Amersham Typhoon™ scanner (Cytiva) with Cy2 or Cy5 filters.

---

**Extended Data Fig. 1:**
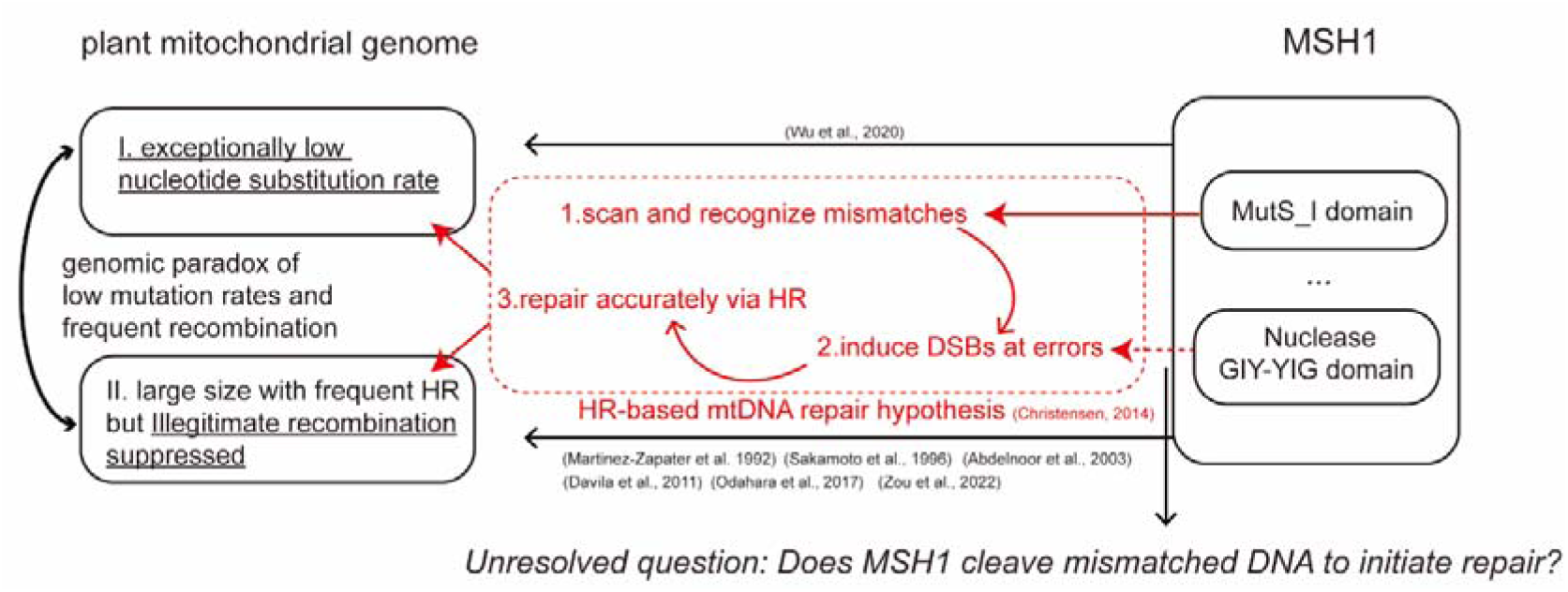
Proposed model of MSH1-dependent plant mitochondrial DNA repair and recombination surveillance in plants. Working hypothesis of plant MSH1-dependent homologous recombination (HR)-based mitochondrial DNA repair (adapted from Christensen, 2014). Most plant mitochondrial genomes are characterized by large size, frequent homologous recombination (HR) but exceptionally low nucleotide substitution rates. MSH1, a nuclear-encoded protein harboring a MutS_I domain and a GIY-YIG endonuclease domain, is proposed to recognize mismatched or improperly paired regions, induce double-strand breaks (DSBs), and facilitate accurate DSB repair through HR, thereby suppressing illegitimate recombination and maintaining genome integrity.

**Extended Data Fig. 2:**
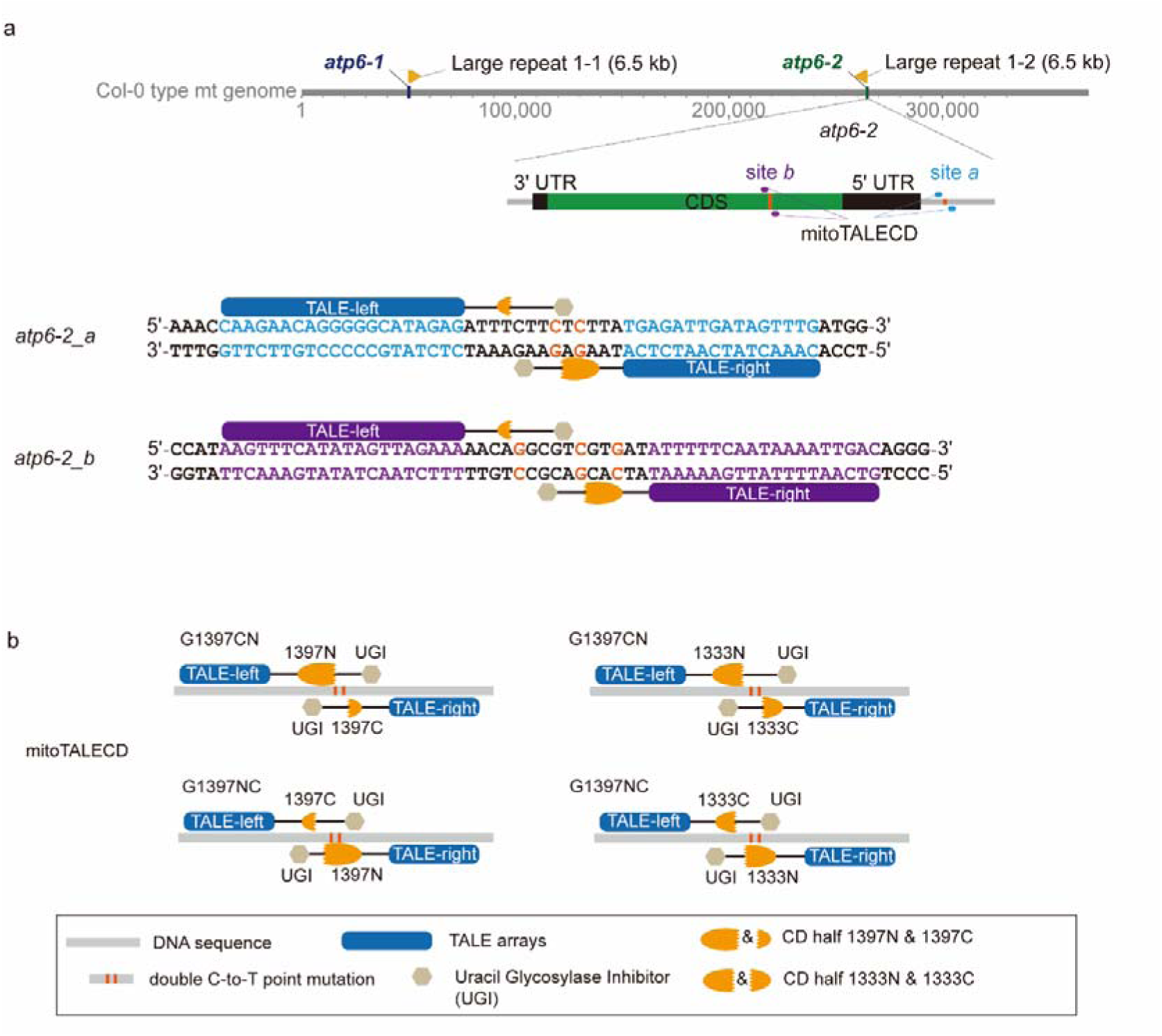
Target site sequences of *atp6-2* and mitoTALECD variants. a, Location of *atp6-2* relative to *atp6-1* and the adjacent large repeats (6.5 kb). Schematic representations of the two target sites, *atp6-2_a* and *atp6-2_b*, with their DNA sequences shown. TALE binding sites are highlighted in blue or purple; the target cytidines for base editing are shown in orange. b, Schematic diagrams of the four mitoTALECD variants (G1397CN, G1397NC, G1333CN, G1333NC) used in this study. TALE arrays, cytidine deaminase halves, and UGI modules are indicated as shown in the legend.

**Extended Data Fig. 3:**
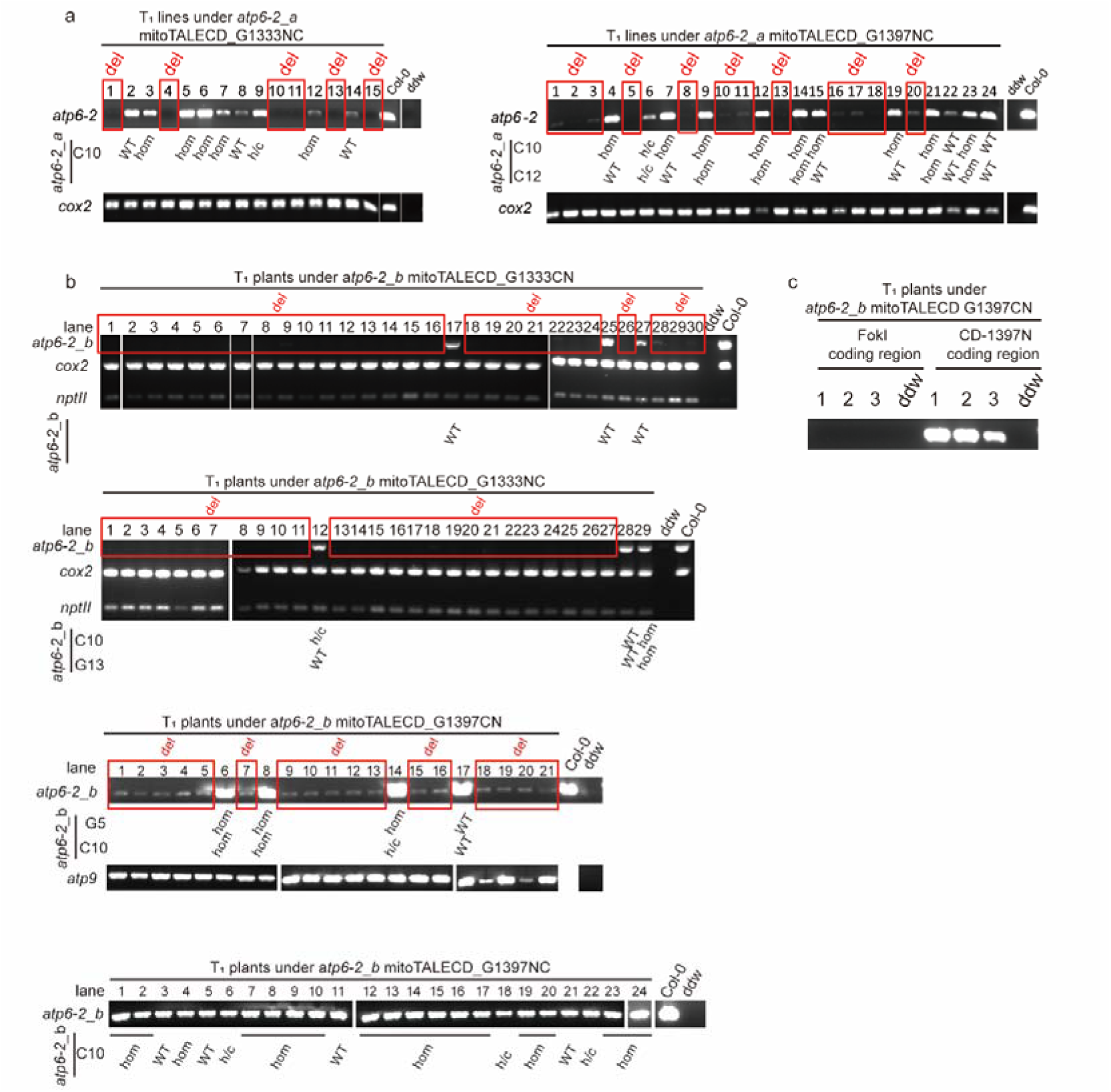
Genotyping T□ plants edited by mitoTALECDs targeting *atp6-2* **a,** PCR and edit results of *atp6-2_a* in T_1_ plants under two types of mitoTALECDs. **b,** PCR and edit results of *atp6-2_b* in T_1_ plants under four types of mitoTALECDs. The numbers following base names (e.g., G5) refer to the position relative to the fifth nucleotide from the left TALE-array binding site (for example, G5 denotes the guanine located five nucleotides downstream of the DNA recognized and bind by left TALE arrays). *nptII*: neomycin phosphotransferase II, a gene that encodes an enzyme that grants resistance to the antibiotic kanamycin in various organisms including plants and bacteria, amplified as reporter gene of mitoTALECD constructs. *cox2* and *atp9* served as a mitochondrial control. Wild type Col-0 was used as a positive control, and ddw was used as a negative control. **c,** PCR of the coding regions of the cytidine deaminase (CD) and FokI domains. Abbreviations: h/c, heteroplasmically or chimaerically substituted; hom, homoplasmically substituted; WT, wild type; ddw, double distilled water.

**Extended Data Fig. 4:**
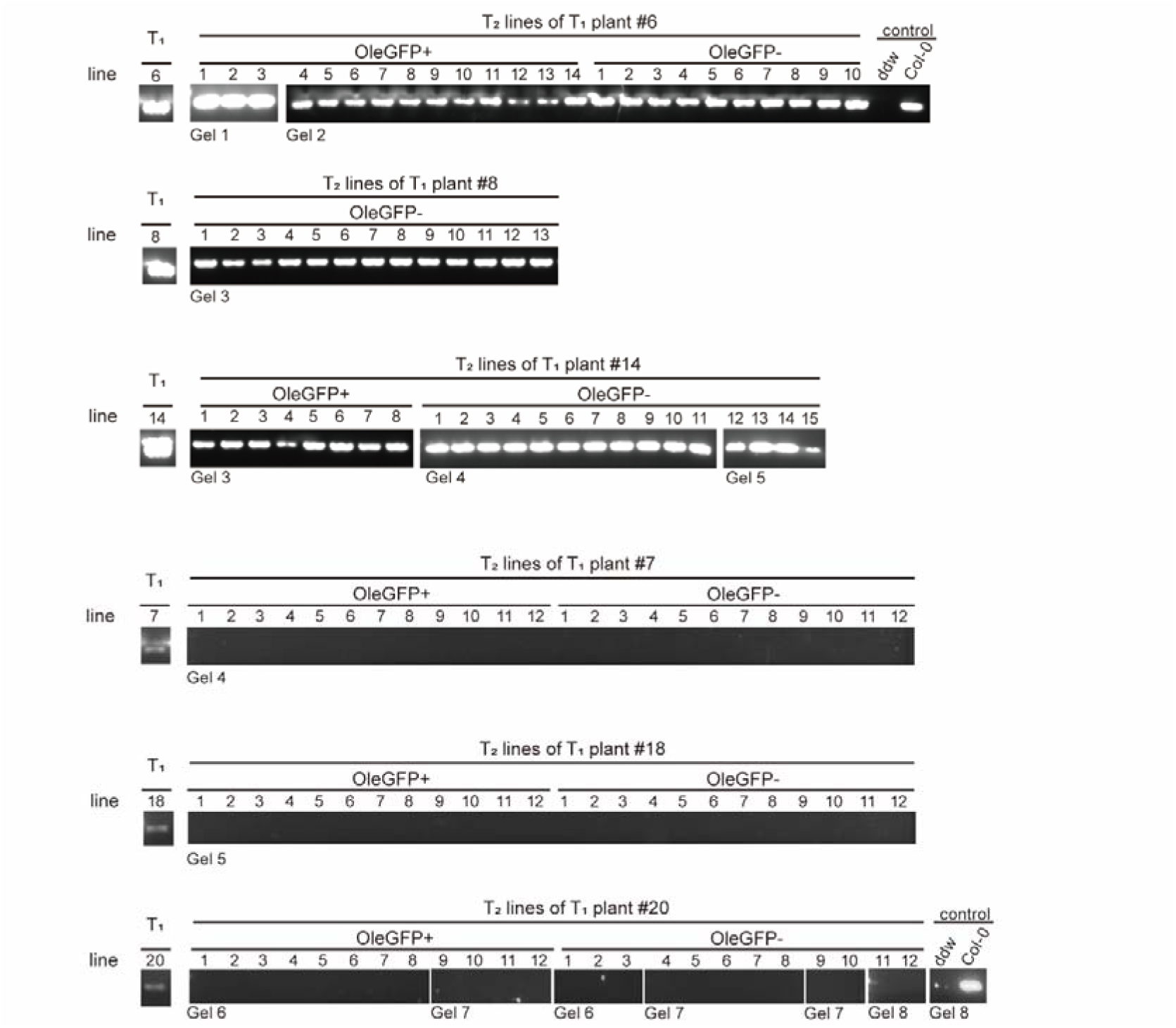
PCR of the *atp6-2_b* in T_2_ progenies derived from six T_1_ lines of *atp6-2_b*_mitoTALECD_G1397CN. Three T_1_ lines with C-to-Tmutation at *atp6-2_b*: #6, #8 and #14; Three T_1_ lines with deletion-like at *atp6-2_b*: #7, #18 and #20. DNA of Col-0 was used as a positive control, and ddw was used as a negative control.

**Extended Data Fig. 5:**
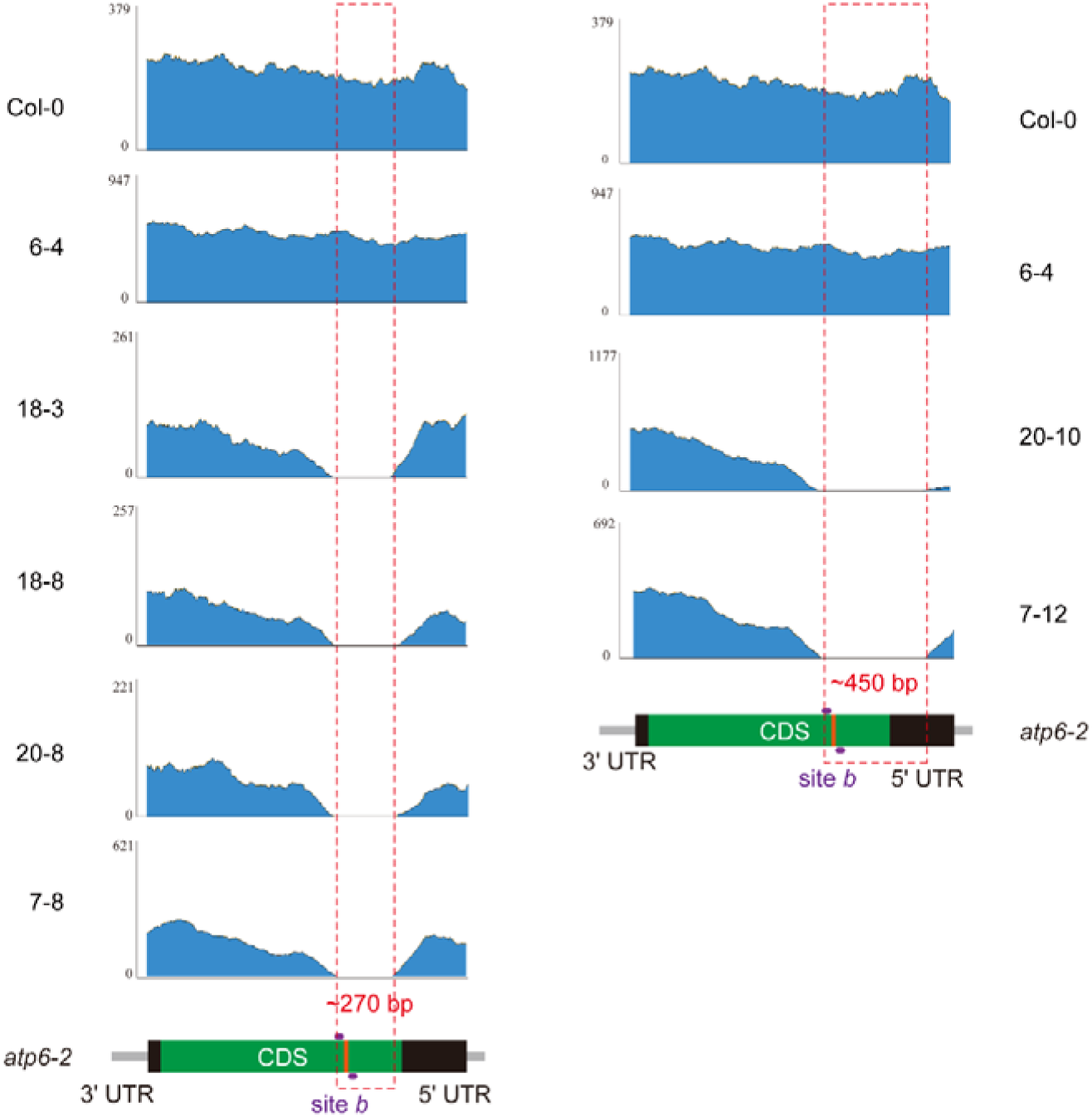
Read mapping at *atp6-2* reveals two distinct deletion patterns in T□ lines edited by *atp6-2_b*_mitoTALECD_G1397CN. Illumina short reads from T□ plants and Col-0 were mapped to the Arabidopsis Col-0 reference mitogenome to assess deletion patterns at the *atp6-2_b*. Coverage plots show sequencing depth across the target region (*atp6-2_b*), with wild-type Col-0 as a control (top). Line 6-4 was derived from a T□ plant that harbored a successful C-to-T conversion at site b but did not exhibit a deletion-like PCR pattern. Lines 18-3, 18-8, 20-8, 7-8, 20-10, and 7-12 were derived from T□ plants previously identified as carrying deletion-like genotypes based on failed PCR amplification at *atp6-2_b*. Among these, four lines (18-3, 18-8, 20-8, 7-8) shared a ∼270-bp deletion, whereas two lines (20-10 and 7-12) exhibited a larger ∼450-bp deletion. Schematic diagrams below the coverage plots llustrate the relative positions and sizes of these deletions at the *atp6-2_b*. The deletions span or flank the base editing target site *b*, which is located within the coding sequence (green bar) of *atp6-2*.

**Extended Data Fig. 6:**
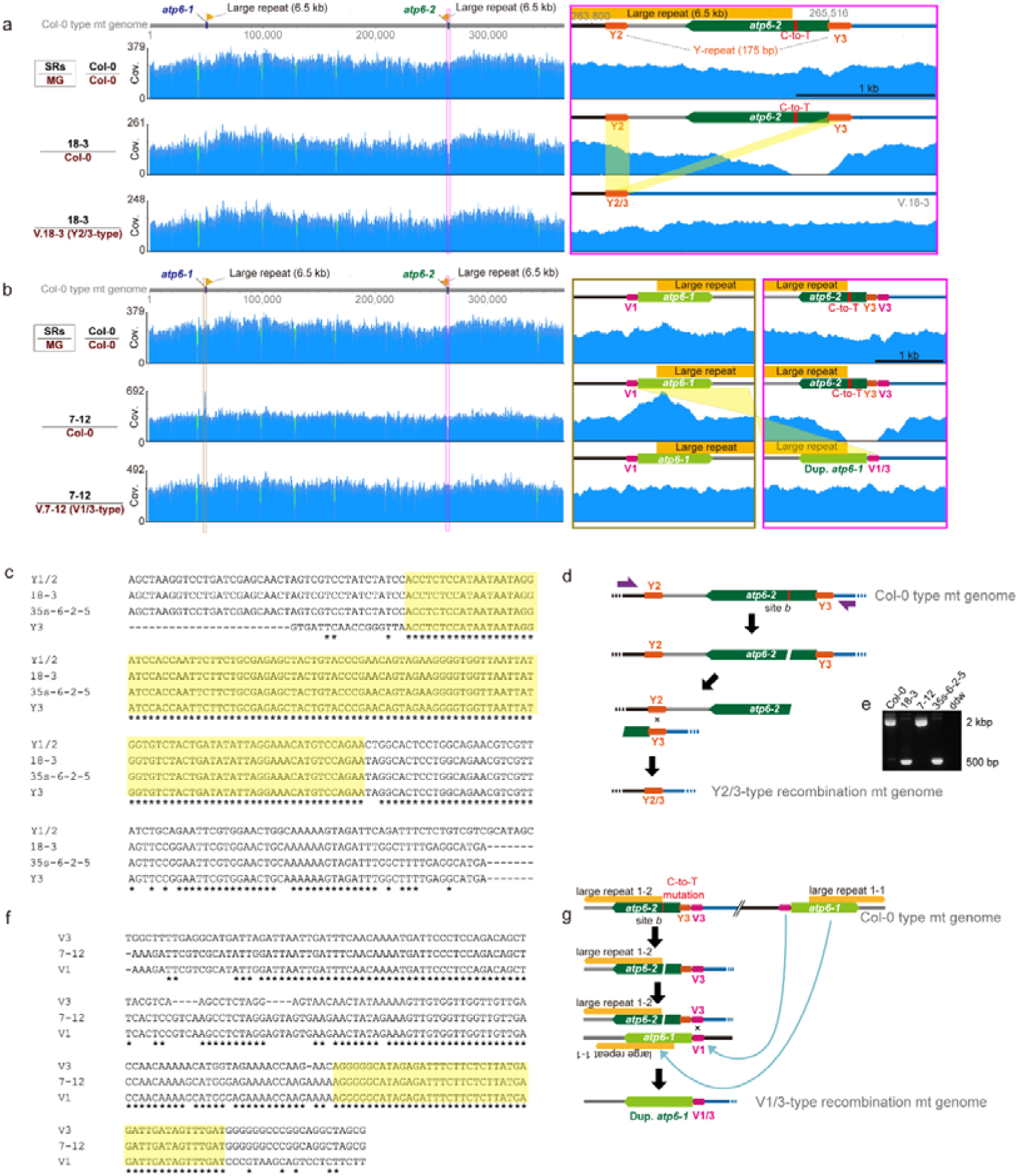
Homoplasmic deletions and two distinct recombination types identified at *atp6-2_b*. **a,b,** Illumina short reads (SRs) from Col-0 and mutant lines 18-3 (a) and 7-12 (b) were mapped to the Col-0 mitochondrial genome (MG; top two rows) and to their respective predicted recombined genomes (bottom row). Green slits on the left indicate apparent deletions in plastid-like regions, which result from the filtering out of short reads derived from the plastid genome. Insets show magnified views of the *atp6* loci. Line 18-3 represents a Y2/3-type recombination event, and line 7-12 represents a V1/3-type recombination event. **c, f,**CLUSTALW alignment of recombined junction sequences in lines 18-3 and 35S-atp6-2-5 (a mitoTALEN-edited line reported in Arimura et al., 2020) with the wild-type Y2/Y3 (c) or V1/V3 (f) repeat regions. Yellow highlights mark identical sequences apparently used in homologous recombination. **d,** Model of homologous recombination between Y2 and Y3 tandem repeats leading to a deletion encompassing *atp6-2_b*. Because of the high similarity between Y1 and Y2, this event is collectively referred to as a Y2/3-type recombination. **e,** PCR detection of Y2/3-type recombination using primers flanking the recombined junction. **g,** Model of the V1/3-type recombination event in line 7-12. The deletion between the V3 repeat and the large repeat near *atp6-2* was repaired by duplication of the region between V1 and the large repeat near *atp6-1*, resulting in a duplicated *atp6-1*.

**Extended Data Fig. 7:**
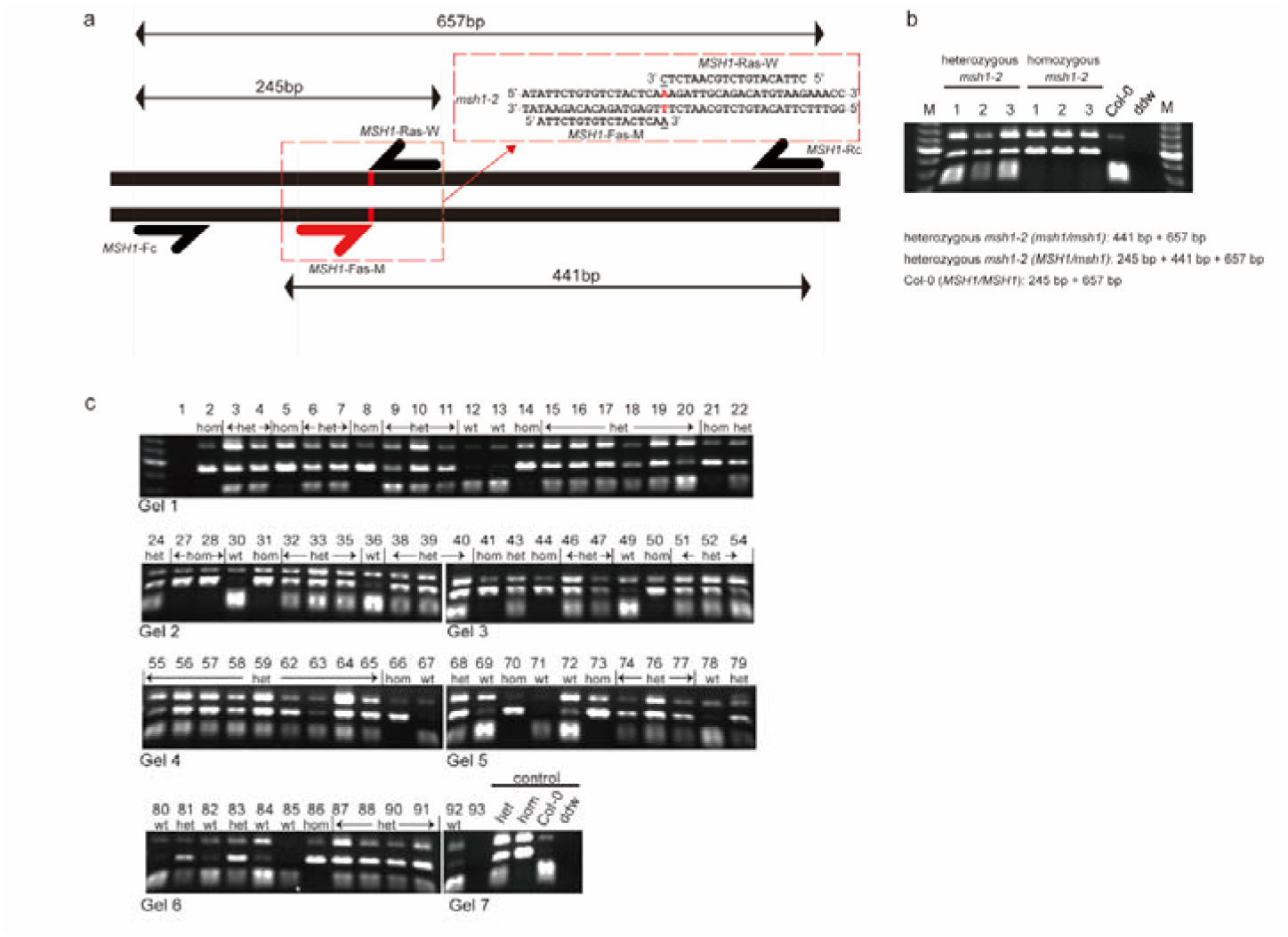
Genotyping strategy and analysis of *msh1-2* alleles in F_2_T_1_ Arabidopsis plants. **a,** Schematic representation of the primer design for allele-specific PCR-based genotyping of the *msh1-2* mutant. A point mutation (marked in red) at the splice site AG before exon 3 of *msh1-2* specifically recognized by allele-specific primers. Primers MSH1-forward allele-specific-mutant (MSH1-Fas-M) and MSH1-reverse allele-specific-WT (MSH1-Ras-W) are designed to selectively amplify the mutant (441 bp) and wild-type (245 bp) alleles, respectively, in combination with the common primers MSH1-Fc and MSH1-Rc. A 657-bp fragment is amplified from both alleles as an internal control. **b,** Validation of allele-specific PCR for *msh1-2* genotyping using Sanger sequencing-confirmed controls. Allele-specific PCR was performed using genomic DNA from three Sanger sequencing-confirmed *msh1-2* homozygous plants (lanes 4–6), three heterozygous plants (lanes 1–3), and Col-0 wild type (lane 7) to validate the specificity and reliability of the primer set. The PCR yields a 657-bp internal control band for all genotypes, a 441-bp band specific to the *msh1-2* mutant allele, a 245-bp band specific to the wild-type allele and both (245-bp and 441-bp) bands in heterogotes. The distinct banding patterns allow discrimination of homozygous, heterozygous, and wild-type genotypes of *msh1-2*. M: DNA ladder. **c,** Genotyping results of F_2_T_1_ transformants. Each lane shows the PCR banding pattern using the primer set in panel a, allowing clear classification of individuals as wild type (wt), heterozygous (het), or homozygous mutant (hom) for *msh1-2*. Control lanes include sequenced genotypes (wt, het, hom) as references, and ddw for double distilled water as negative control.

**Extended Data Fig. 8:**
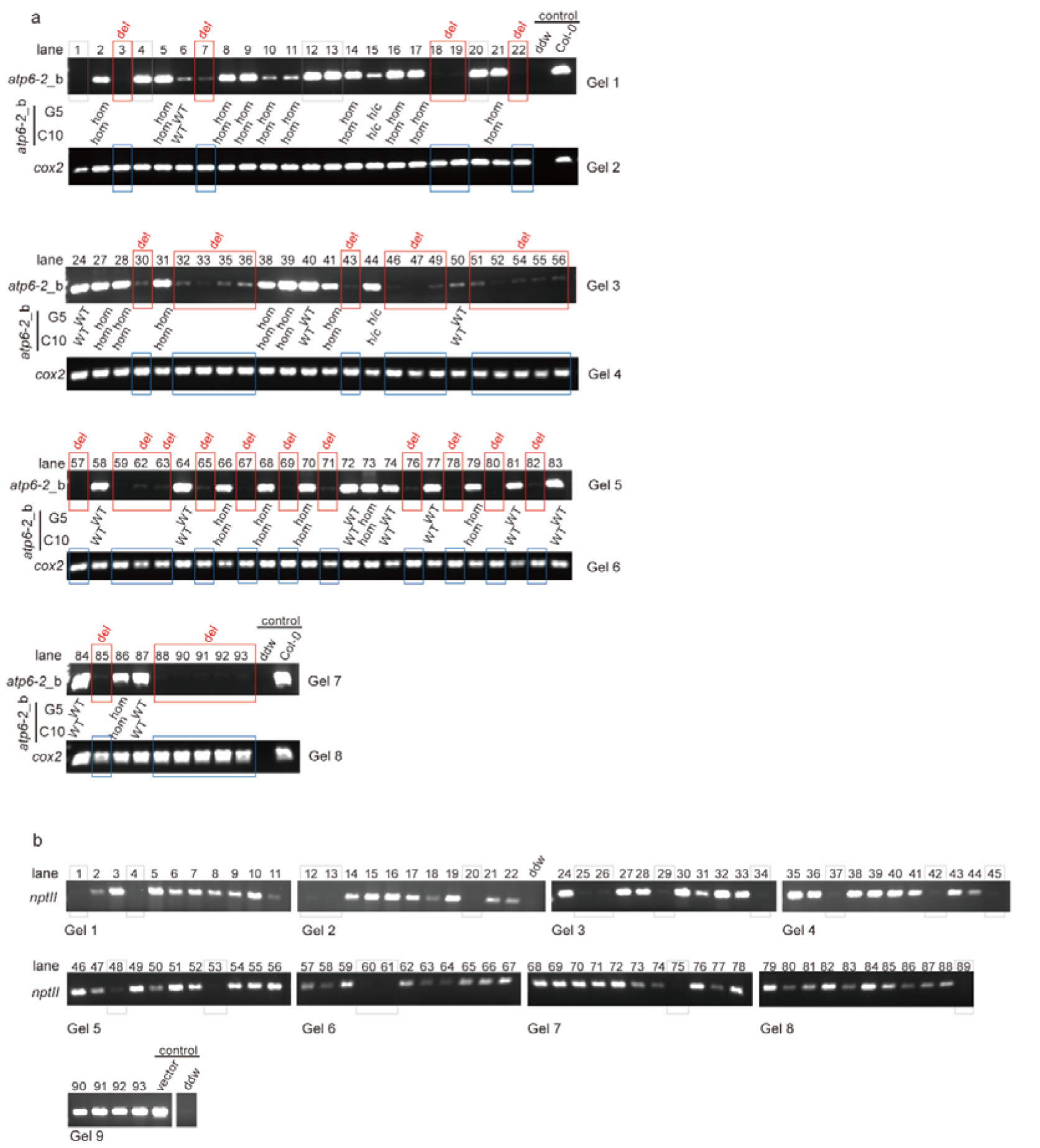
Genotyping *atp6-2*_*b* of F_2_T_1_ Arabidopsis plants. **a**, Genotyping *atp6-2_b* of F_2_T_1_ plants under *atp6-2_b*_mitoTALECD_G1397CN. F□T□ lines that failed to amplify at the *atp6-2*_*b* were identified as deletion mutants (del) and indicated by red boxes. The same lines were successfully amplified at *cox2*, which serves as a positive control, and are marked by blue boxes. F□T□ lines that were successfully amplified at *atp6-2*_b were further analyzed by Sanger sequencing. G5 and C10 represent editing window positions corresponding to the fifth guanine (G5) and tenth cytosine (C10) nucleotides downstream from the left TALE array binding sequence. Wild-type (WT) denotes lines with no mutations detected. “Hom” indicates homoplasmic C-to-T mutations at the given base, whereas “h/c” indicates heteroplasmic or chimeric C-to-T mutations. Lines with invalid DNA or identified as non-transformants based on *nptII* PCR were excluded from the figure or marked with grey boxes and omitted from the final analysis. Double distilled water and wild type Col-0 templates were conducted as negative and positive control, respectively. **b,** *nptII* PCR validation for all F□T□ lines. Lines that failed to amplify *nptII* are marked with grey boxes and were not included in subsequent statistical analyses. A 1:1,000 diluted *atp6-2_b*_mitoTALECD_G1397CN vector was used as a positive control, and ddw was used as a negative control.

**Extended Data Fig. 9:**
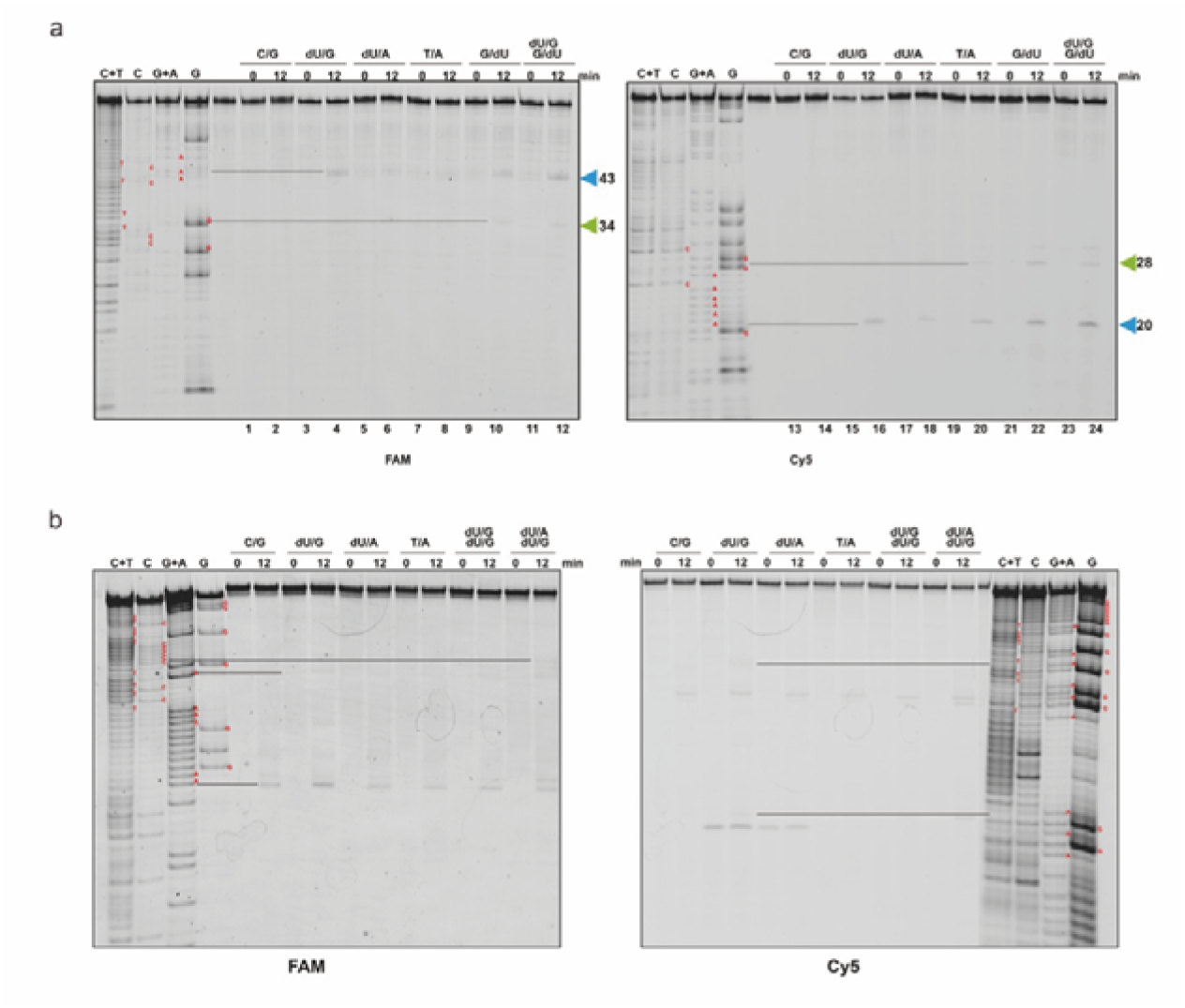
Mapping of MSH1 cleavage sites by Maxam–Gilbert sequencing of *atp6-2_b* and *atp6-2_a*. **a,** Mapping of MSH1 cleavage sites by Maxam–Gilbert sequencing of *atp6-2_b.* **b,** Mapping of MSH1 cleavage sites by Maxam–Gilbert sequencing of *atp6-2_a.* 12% denaturing gel showing MSH1 cleavage reactions side by side with Maxam–Gilbert sequencing ladders of the 5′-FAM- and 5′-Cy5-labeled oligonucleotides used to assemble the wild-type (C:G) *atp6-2_*b sequence. Cleavage sites are indicated by lines and inferred from the ladders and are mapped onto the corresponding DNA sequence using cyan and green triangles.

**Extended Data Fig. 10:**
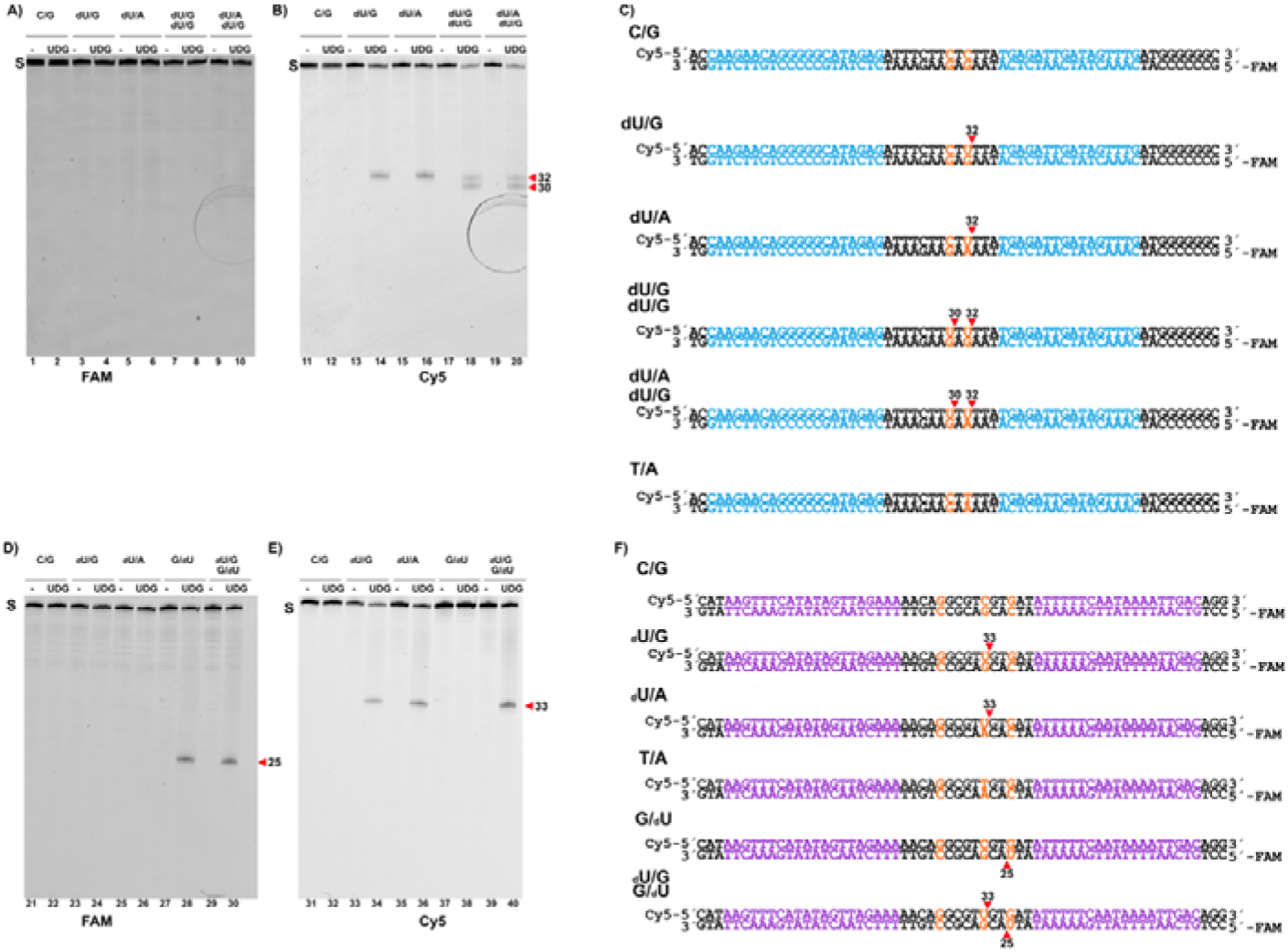
Identification of uracil-containing DNA substrates by UDG cleavage on a denaturing urea gel. **a to e**, dsDNA substrates incubated in the presence (+UDG) or absence (–UDG) of uracil-DNA glycosylase (UDG) and analyzed by denaturing polyacrylamide gel electrophoresis containing urea. Excision of uracil by UDG generates abasic (AP) sites that undergo strand cleavage under the presence of NaOH and denaturing conditions, DNA fragments that indicate the presence of uracil. Fluorescently labeled substrates containing uracil display a clear cleavage from full-length species to defined cleavage products upon UDG treatment, whereas substrates lacking uracil remain intact. Separation on the urea gel enables the visualization of cleavage products, confirming both the presence and position of uracil within the substrate. dsDNA oligonucleotide sequences and fragment sizes (nt) are shown on the left (**c and f**).

## Notes

### Competing Interest Statement

The authors have declared no competing interest.

### Funding

- JSPS: 24H02271, JSCCA20230008

## REFERENCES

Abdelnoor, R. V., A. C. Christensen, S. Mohammed, B. Munoz-Castillo, H. Moriyama and S. A. Mackenzie (2006). “Mitochondrial genome dynamics in plants and animals: convergent gene fusions of a MutS homologue.” J Mol Evol 63(2): 165–173.

Abdelnoor, R. V., R. Yule, A. Elo, A. C. Christensen, G. Meyer-Gauen and S. A. Mackenzie (2003). Substoichiometric shifting in the plant mitochondrial genome is influenced by a gene homologous to MutS. Proc Natl Acad Sci U S A. United States. 100: 5968–5973.

Arimura, S., J. Yamamoto, G. P. Aida, M. Nakazono and N. Tsutsumi (2004). “Frequent fusion and fission of plant mitochondria with unequal nucleoid distribution.” Proc Natl Acad Sci U S A 101(20): 7805–7808.

Arimura, S. I. (2018). “Fission and Fusion of Plant Mitochondria, and Genome Maintenance.” Plant Physiol 176(1): 152–161.

Arimura, S. I., H. Ayabe, H. Sugaya, M. Okuno, Y. Tamura, Y. Tsuruta, Y. Watari, S. Yanase, T. Yamauchi, T. Itoh, A. Toyoda, H. Takanashi and N. Tsutsumi (2020). “Targeted gene disruption of ATP synthases 6-1 and 6-2 in the mitochondrial genome of Arabidopsis thaliana by mitoTALENs.” Plant J 104(6): 1459–1471.

Ayala-Garcia, V. M., N. Baruch-Torres, P. L. Garcia-Medel and L. G. Brieba (2018). “Plant organellar DNA polymerases paralogs exhibit dissimilar nucleotide incorporation fidelity.” FEBS J 285(21): 4005–4018.

Broz, A. K., M. M. Hodous, Y. Zou, P. C. Vail, Z. Wu and D. B. Sloan (2025). Flipping the switch on some of the slowest mutating genomes: Direct measurements of plant mitochondrial and plastid mutation rates in msh1 mutants. PLoS Genet. United States. 21: e1011764.

Broz, A. K., A. Keene, M. Fernandes Gyorfy, M. Hodous, I. G. Johnston and D. B. Sloan (2022). “Sorting of mitochondrial and plastid heteroplasmy in Arabidopsis is extremely rapid and depends on MSH1 activity.” Proc Natl Acad Sci U S A 119(34): e2206973119.

Chevigny, N., D. Schatz-Daas, F. Lotfi and J. M. Gualberto (2020). “DNA Repair and the Stability of the Plant Mitochondrial Genome.” Int J Mol Sci 21(1).

Christensen, A. C. (2013). “Plant mitochondrial genome evolution can be explained by DNA repair mechanisms.” Genome Biol Evol 5(6): 1079–1086.

Christensen, A. C. (2014). “Genes and Junk in Plant Mitochondria-Repair Mechanisms and Selection.” Genome Biology and Evolution 6(6): 1448–1453.

Clough, S. J. and A. F. Bent (1998). “Floral dip: a simplified method for Agrobacterium-mediated transformation of Arabidopsis thaliana.” Plant J 16(6): 735–743.

Coulon, S., P. H. L. Gaillard, C. Chahwan, W. H. McDonald, J. R. Yates, 3rd and P. Russell (2004). “Slx1-Slx4 are subunits of a structure-specific endonuclease that maintains ribosomal DNA in fission yeast.” Mol Biol Cell 15(1): 71–80.

Davila, J. I., M. P. Arrieta-Montiel, Y. Wamboldt, J. Cao, J. Hagmann, V. Shedge, Y. Z. Xu, D. Weigel and S. A. Mackenzie (2011). “Double-strand break repair processes drive evolution of the mitochondrial genome in Arabidopsis.” BMC Biol 9: 64.

Dryden, D. T., Murray, N. E. & Rao, D. N. Nucleoside triphosphate-dependent restriction enzymes. Nucleic Acids Res.29, 3728–3741 (2001).

Dunin-Horkawicz, S., M. Feder and J. M. Bujnicki (2006). “Phylogenomic analysis of the GIY-YIG nuclease superfamily.” Bmc Genomics 7.

Edgell, D. R., V. Derbyshire, P. Van Roey, S. LaBonne, M. J. Stanger, Z. Li, T. M. Boyd, D. A. Shub and M. Belfort (2004). “Intron-encoded homing endonuclease I-TevI also functions as a transcriptional autorepressor.” Nat Struct Mol Biol 11(10): 936–944.

Fields, P. D., G. Waneka, M. Naish, M. C. Schatz, I. R. Henderson and D. B. Sloan (2022). Complete Sequence of a 641-kb Insertion of Mitochondrial DNA in the Arabidopsis thaliana Nuclear Genome. Genome Biol Evol. England. 14.

Fukui, K., A. Harada, T. Wakamatsu, A. Minobe, K. Ohshita, M. Ashiuchi and T. Yano (2018). “The GIY-YIG endonuclease domain of Arabidopsis MutS homolog 1 specifically binds to branched DNA structures.” FEBS Lett 592(24): 4066–4077.

Gandini, C. L., L. E. Garcia, C. C. Abbona, L. F. Ceriotti, S. Kushnir, D. Geelen and M. V. Sanchez-Puerta (2023). “Break-induced replication is the primary recombination pathway in plant somatic hybrid mitochondria: a model for mitochondrial horizontal gene transfer.” J Exp Bot 74(12): 3503–3517.

Gualberto, J. M., D. Mileshina, C. Wallet, A. K. Niazi, F. Weber-Lotfi and A. Dietrich (2014). “The plant mitochondrial genome: dynamics and maintenance.” Biochimie 100: 107–120.

Gualberto, J. M. and K. J. Newton (2017). “Plant Mitochondrial Genomes: Dynamics and Mechanisms of Mutation.” Annu Rev Plant Biol 68: 225–252.

Kang, B. C., S. J. Bae, S. Lee, J. S. Lee, A. Kim, H. Lee, G. Baek, H. Seo, J. Kim and J. S. Kim (2021). “Chloroplast and mitochondrial DNA editing in plants.” Nat Plants 7(7): 899–905.

Kazama, T., M. Okuno, Y. Watari, S. Yanase, C. Koizuka, Y. Tsuruta, H. Sugaya, A. Toyoda, T. Itoh, N. Tsutsumi, K. Toriyama, N. Koizuka and S. I. Arimura (2019). “Curing cytoplasmic male sterility via TALEN-mediated mitochondrial genome editing.” Nat Plants 5(7): 722–730.

Kisker, C., J. Kuper and B. Van Houten (2013). “Prokaryotic nucleotide excision repair.” Cold Spring Harb Perspect Biol 5(3): a012591.

Kubo, T. and K. J. Newton (2008). “Angiosperm mitochondrial genomes and mutations.” Mitochondrion 8(1): 5–14.

Kunkel, T. A. and D. A. Erie (2015). “Eukaryotic Mismatch Repair in Relation to DNA Replication.” Annu Rev Genet 49: 291–313.

Larson, E. D., D. W. Bednarski and N. Maizels (2008). “High-fidelity correction of genomic uracil by human mismatch repair activities.” BMC Mol Biol 9: 94.

Lencina, F., A. R. Prina, M. G. Pacheco, K. Kobayashi and A. M. Landau (2025). Four Large Indels in Barley Chloroplast Mutator (cpm) Seedlings Reinforce the Hypothesis of a Malfunction in the MMR System. Int J Mol Sci. Switzerland. 26.

Lin, Z., M. Nei and H. Ma (2007). “The origins and early evolution of DNA mismatch repair genes--multiple horizontal gene transfers and co-evolution.” Nucleic Acids Res 35(22): 7591–7603.

Martinez-Zapater, J. M., P. Gil, J. Capel and C. R. Somerville (1992). “Mutations at the Arabidopsis CHM locus promote rearrangements of the mitochondrial genome.” Plant Cell 4(8): 889–899.

Mazurek, A., M. Berardini and R. Fishel (2002). “Activation of human MutS homologs by 8-oxo-guanine DNA damage.” J Biol Chem 277(10): 8260–8266.

McCulloch, S. D. and T. A. Kunkel (2008). “The fidelity of DNA synthesis by eukaryotic replicative and translesion synthesis polymerases.” Cell Res 18(1): 148–161.

Modrich, P. and R. Lahue (1996). “Mismatch repair in replication fidelity, genetic recombination, and cancer biology.” Annu Rev Biochem 65: 101–133.

Mok, B. Y., M. H. de Moraes, J. Zeng, D. E. Bosch, A. V. Kotrys, A. Raguram, F. Hsu, M. C. Radey, S. B. Peterson, V. K. Mootha, J. D. Mougous and D. R. Liu (2020). “A bacterial cytidine deaminase toxin enables CRISPR-free mitochondrial base editing.” Nature 583(7817): 631–637.

Morley, S. A. and B. L. Nielsen (2017). “Plant mitochondrial DNA.” Front Biosci (Landmark Ed) 22(6): 1023–1032.

Mueller, J. E., D. Smith, M. Bryk and M. Belfort (1995). “Intron-encoded endonuclease I-TevI binds as a monomer to effect sequential cleavage via conformational changes in the td homing site.” EMBO J 14(22): 5724–5735.

Nakazato, I., M. Okuno, H. Yamamoto, Y. Tamura, T. Itoh, T. Shikanai, H. Takanashi, N. Tsutsumi and S. I. Arimura (2021). “Targeted base editing in the plastid genome of Arabidopsis thaliana.” Nat Plants 7(7): 906–913.

Nakazato, I., M. Okuno, C. Zhou, T. Itoh, N. Tsutsumi, M. Takenaka and S. I. Arimura (2022). “Targeted base editing in the mitochondrial genome of Arabidopsis thaliana.” Proc Natl Acad Sci U S A 119(20): e2121177119.

Odahara, M. (2020). “Factors Affecting Organelle Genome Stability in Physcomitrella patens.” Plants (Basel) 9(2).

Odahara, M. and Y. Sekine (2018). “RECX Interacts with Mitochondrial RECA to Maintain Mitochondrial Genome Stability.” Plant Physiol 177(1): 300–310.

Ogata, H., J. Ray, K. Toyoda, R. A. Sandaa, K. Nagasaki, G. Bratbak and J. M. Claverie (2011). “Two new subfamilies of DNA mismatch repair proteins (MutS) specifically abundant in the marine environment.” ISME J 5(7): 1143–1151.

On, Y. Y. and M. Welch (2021). “The methylation-independent mismatch repair machinery in Pseudomonas aeruginosa.” Microbiology (Reading) 167(12).

Paszkiewicz, G., J. M. Gualberto, A. Benamar, D. Macherel and D. C. Logan (2017). “Arabidopsis Seed Mitochondria Are Bioenergetically Active Immediately upon Imbibition and Specialize via Biogenesis in Preparation for Autotrophic Growth.” Plant Cell 29(1): 109–128.

Penafiel-Ayala, A., A. Peralta-Castro, J. Mora-Garduno, P. Garcia-Medel, A. G. Zambrano-Pereira, C. Diaz-Quezada, M. J. Abraham-Juarez, C. G. Benitez-Cardoza, D. B. Sloan and L. G. Brieba (2024). “Plant Organellar MSH1 Is a Displacement Loop-Specific Endonuclease.” Plant Cell Physiol 65(4): 560–575.

Peñafiel-Ayala, A., C. Zhou, D. B. Sloan, S.-i. Arimura and L. G. Brieba (2026). “Plant MutS Homolog 1 is a mismatch-directed nuclease required for organelle genome maintenance” https://www.biorxiv.org/.

Preuten, T., E. Cincu, J. Fuchs, R. Zoschke, K. Liere and T. Borner (2010). “Fewer genes than organelles: extremely low and variable gene copy numbers in mitochondria of somatic plant cells.” Plant J 64(6): 948–959.

Sakuma, T. and T. Yamamoto (2016). “Engineering Customized TALENs Using the Platinum Gate TALEN Kit.” Methods Mol Biol 1338: 61–70.

Shimada, T. L., T. Shimada and I. Hara-Nishimura (2010). “A rapid and non-destructive screenable marker, FAST, for identifying transformed seeds of Arabidopsis thaliana.” Plant J 61(3): 519–528.

Sloan, D. B., A. K. Broz, S. A. Kuster, V. Muthye, A. Penafiel-Ayala, J. R. Marron, D. V. Lavrov and L. G. Brieba (2025). “Expansion of the MutS gene family in plants.” Plant Cell 37(7).

Tamburino, R., N. D’Agostino, G. Aufiero, A. Nicolia, A. Facchiano, D. Giordano, L. Sannino, R. Paparo, S. I. Arimura, N. Scotti and T. Cardi (2025). “Mitochondrial gene editing and allotopic expression unveil the role of orf125 in the induction of male fertility in some Solanum spp. hybrids and in the evolution of the common potato.” Plant Biotechnol J 23(5): 1862–1875.

Wang, D. Y., Q. Zhang, Y. Liu, Z. F. Lin, S. X. Zhang, M. X. Sun and Sodmergen (2010). “The levels of male gametic mitochondrial DNA are highly regulated in angiosperms with regard to mitochondrial inheritance.” Plant Cell 22(7): 2402–2416.

Wu, Z., G. Waneka, A. K. Broz, C. R. King and D. B. Sloan (2020). “MSH1 is required for maintenance of the low mutation rates in plant mitochondrial and plastid genomes.” Proc Natl Acad Sci U S A 117(28): 16448–16455.

Zhou, C., M. Okuno, I. Nakazato, N. Tsutsumi and S. I. Arimura (2024). “Targeted A-to-G base editing in the organellar genomes of Arabidopsis with monomeric programmable deaminases.” Plant Physiol 194(4): 2278–2287.

Zou, Y., W. Zhu, D. B. Sloan and Z. Wu (2022). “Long-read sequencing characterizes mitochondrial and plastid genome variants in Arabidopsis msh1 mutants.” Plant J 112(3): 738–755.

